# Visualizing sub-organellar lipid distribution using correlative light and electron microscopy

**DOI:** 10.1101/2025.01.28.635245

**Authors:** H. Mathilda Lennartz, Suman Khan, Kristin Böhlig, Weihua Leng, Falk Elsner, Nadav Scher, Michaela Wilsch-Bräuninger, Ori Avinoam, André Nadler

## Abstract

Lipids and proteins compartmentalize biological membranes into nanoscale domains which are crucial for signaling, intracellular trafficking and many other cellular processes. Studying nanodomain function requires the ability to measure protein and lipid localization at the nanoscale. Current methods for visualizing lipid localization do not meet this requirement. Here, we introduce a correlative light and electron microscopy workflow to image lipids (Lipid-CLEM), combining near-native lipid probes and on-section labeling by click chemistry. This approach enables the quantification of relative lipid densities in membrane nanodomains. We find differential partitioning of sphingomyelin into intraluminal vesicles, recycling tubules, and the boundary membrane of the early endosome, representing a degree of nanoscale organization previously observed only for proteins. We anticipate that our Lipid-CLEM workflow will greatly facilitate the mechanistic analysis of lipid functions in cell biology, allowing for the simultaneous investigation of proteins and lipids during membrane nanodomain assembly and function.

## Introduction

The eucaryotic cell is compartmentalized by membranes, which are organized into functional nano- and mesoscale domains that feature distinct geometries, lifetimes, and protein compositions. Membrane domains have been implicated in cell signaling ^1^, cell adhesion ^1,2^, membrane remodeling ^3^ and trafficking ^3,4^. In contrast to the well-studied protein composition of membrane domains, much less is known about their lipid content, as visualizing individual lipid species at the ultrastructural level is largely an unsolved methodological problem ^5,6^. Key challenges include rapid lipid dynamics ^7^ and localization artifacts introduced by fluorescently labeled lipid probes ^8–10^. A suitable lipid visualization technique should report on molecularly distinct lipid species rather than entire lipid classes and employ minimally modified lipid probes to avoid artifacts. Furthermore, membrane ultrastructure must be preserved and spatially resolved. Finally, high temporal resolution during sample preparation must be achieved to capture dynamic membrane remodeling events.

Bifunctional diazirine-alkyne lipid probes (“bifunctional lipids” in the following) are a powerful tool to visualize lipid localization by fluorescence microscopy *in situ* ^11–16^. Bifunctional lipids closely mimic native lipid species as their small chemical modifications limit localization artifacts to a minimum ^17,18^. They can be introduced into membranes of living cells and rapidly photo-crosslinked to neighboring proteins ^19^. The resulting lipid-protein conjugates can, in turn, be fluorescently labeled post-fixation by click chemistry, circumventing the use of heavily modified lipid probes in living cells while capturing lipids that cannot be chemically fixed otherwise ^18,20^. Fluorescence lipid imaging based on bifunctional probes has been instrumental for mapping inter-organelle lipid distribution and lipid transport pathways ^20,21^.

Measuring lipid localization at the nanoscale, however, requires a resolution that light microscopy alone cannot provide ^22^. A correlative light and electron microscopy (CLEM) approach ^23^ to measure lipid densities within nanodomains would allow us to investigate lipid localization-function relationships ^24^, e.g. regarding lipid sorting mechanisms during membrane trafficking ^25^. CLEM approaches to image lipids have been reported previously by us^12^ and others ^26,27^. The required sample processing procedures of these approaches, however, are typically not ideal for the preservation of membrane ultrastructure, are limited to studying the outer leaflet of the plasma membrane, or target entire lipid classes.

Here we report a Lipid-CLEM workflow based on bifunctional lipid probes for faithfully visualizing lipids on the ultrastructural level. Our CLEM workflow is optimized for membrane ultrastructure preservation, sample preparation speed, and correlation precision. We used a sequence of rapid lipid photo-crosslinking ^19^ and high-pressure freezing ^28,29^ to arrest lipid localization within the sample, followed by freeze substitution ^28^, sectioning, and on-section fluorescence labeling of lipid probes by click chemistry. The combined time required for crosslinking and high-pressure freezing (approximately 10 s) allows for capturing membrane domains with lifetimes in the second to minute range. We developed Lipid-CLEM variants that either feature labeled bifunctional lipids solely on the surface of the section (optimized for membrane ultrastructure preservation) or throughout the section (optimized for lipid density measurements).

Using whole-section labeling, we investigated the capacity of the early endosome to sort lipids. It is well established that proteins within the early endosome segregate into distinct membrane domains as a prerequisite for sorting into different trafficking pathways ^30–33^. For instance, transferrin segregates into recycling tubules to be trafficked back to the plasma membrane ^31,33^ and low-density lipoprotein particles (LDL) accumulate in the endosomal lumen for transport towards lysosomes ^32,33^. Whether lipids are sorted in a similar fashion during retrograde membrane trafficking is not known ^25^. Several studies have addressed this question indirectly by quantifying uptake of lipid-fluorophore conjugates into endosomes ^34,35^, but due to the experimental setup, direct partitioning into endosomal sub-compartments could not be assessed. Using Lipid-CLEM we measured the density of a single sphingomyelin species in the sub-compartments of individual early endosomes. We found that sphingomyelin is relatively enriched in intraluminal vesicles and depleted from recycling tubules, suggesting active lipid sorting mechanisms within the early endosome. Furthermore, protein and lipid cargoes exhibit distinctly different localization patterns within the endosomal membrane system, implying that lipid and protein cargo trafficking routes diverge in the early endosome.

## Results

### Bifunctional lipid probes enable on-section Lipid-CLEM

In principle, Lipid-CLEM can be performed using cryo- or room-temperature (RT) workflows. Cryo-CLEM facilitates the highest membrane preservation but necessitates the use of fluorescently labeled lipids in the living cell. On the other hand, RT-CLEM allows for the use of near-native bifunctional lipid probes while compromising on ultrastructural preservation (Extended Data 1A). To assess the viability of both strategies, we first compared two phosphatidylcholine (PC) derivatives, a fluorescent nitrobenzoxadiazole (NBD)-PC and its near-native bifunctional PC analog (Extended Data 1 B). We found that the transport behavior of the fluorescent NBD-PC deviated significantly from the bifunctional lipid probe (Extended Data 1 C-E). This implies a tradeoff between the improved ultrastructure of cryo-preserved samples using fluorescent lipid analogs and much more faithful recapitulation of native lipid behavior by minimally modified lipid probes. We thus opted for a RT Lipid-CLEM approach (Figure 1 A, B) using bifunctional lipids based on our recently reported workflow for fluorescence imaging of lipids ^20^.

**Figure 1.**
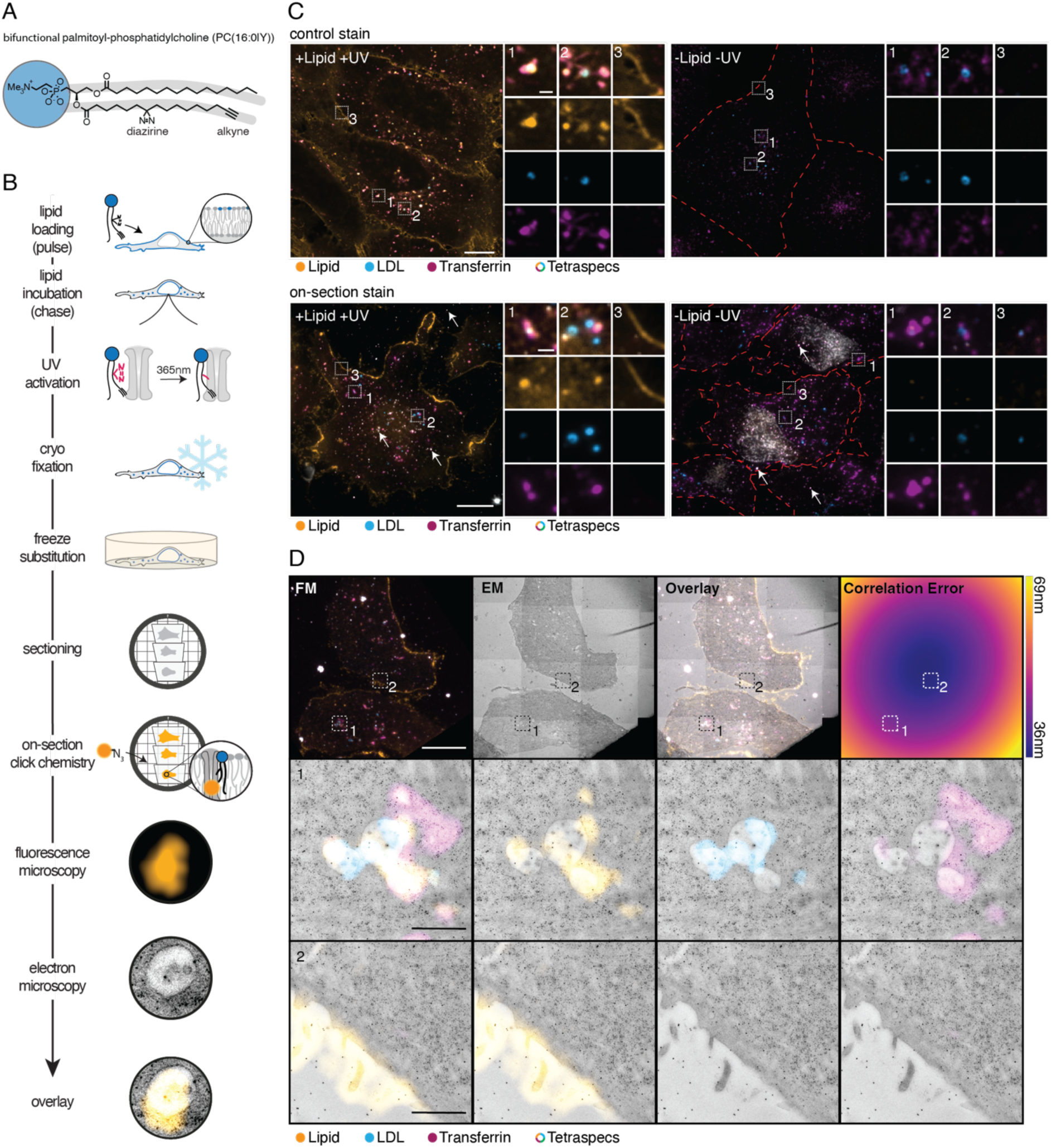
On-section copper catalyzed click reaction for visualizing lipids using CLEM. A: Chemical structure of the bifunctional lipid probe PC(16:0|Y). B: Lipid-CLEM workflow: Cells are loaded with bifunctional lipid probes, probes are UV-crosslinked and samples are high-pressure frozen. Lipids are fluorescently labelled directly on sections after resin-embedding via freeze-substitution. Samples are imaged by fluorescence and electron microscopy. Images are overlayed by correlation. C: Cells were co-labelled with fluorescent LDL (blue) and transferrin (magenta). Lipid signal is shown in orange. Control stain: Cells with (+Lipid +UV) and without (-Lipid -UV) bifunctional PC(16:0|Y) were chemically fixed, permeabilized, labelled by click chemistry and imaged using confocal fluorescence microscopy. On-section stain: Thin sections (100nm) of HM20 samples treated with (+Lipid +UV) and without (-Lipid -UV) bifunctional PC(16:0|Y) were labelled according to the workflow. Multicolored Tetrapecs are indicated by arrows. Dotted red lines represent cell borders if not visible otherwise. Scale bar of overview images: 10 µm; scale bar of ROIs: 1 µm. D: CLEM of thin HM20 sections (100nm) shows localization of the lipid at the plasma membrane and endosomes. The fluorescent overview image (FM), medium magnified TEM image (EM), overlay and correlation error map of 100 nm thick sections are shown left to right. Representative ROIs for an endosome (1) and the plasma membrane (2) are shown, acquired at higher magnification by TEM. Scale bar of overview images: 10 µm; Scale bar of ROIs: 1 µm.

We aimed to maximize temporal resolution, preservation of membrane ultrastructure and lipid signal correlation precision. Thus, we decided to introduce the fluorescent label directly on the section as a final step of the sample processing routine and omitted initial chemical permeabilization steps to benefit from the higher temporal resolution of high pressure freezing. This resulted in the following workflow: Bifunctional lipid probes were loaded into the outer leaflet of the plasma membrane of living cells using an alpha-methyl-cyclodextrin-mediated lipid exchange reaction between donor liposomes and the plasma membrane (Extended Data 2). After a 4 min chase time, lipid probes were photo-crosslinked to generate covalent lipid-protein conjugates, followed by immediate cryo-fixation using high-pressure freezing (Figure 1 B). For CLEM experiments, cells were cultured on carbon-coated sapphire disks. We found that the carbon coat reduced photoactivation efficiency by absorption of UV light (Extended Data 3). To improve lipid crosslinking, we developed a 365 nm (±10 nm) UV-LED device, which was placed directly above the sample in the aqueous medium. A spacer ring between the sample and the LED ensured homogenous illumination. With this setup, a 3 s irradiation pulse was sufficient to achieve signal intensities comparable to irradiation through uncoated coverslips (Extended Data 3). Crosslinked and frozen samples were subjected to freeze substitution to dehydrate and embed samples into resin. This procedure also removed residual bifunctional lipid probes not crosslinked to proteins (Extended Data 4), which greatly reduced unspecific fluorescence lipid signal. Resin-embedded samples were sectioned, and individual sections were stained by on-section copper-catalyzed click chemistry (“click chemistry” in the following) ^36,37^. Sections were stained twice since we observed an increase in signal-to-noise ratio after repeated click labeling (Extended Data 5). Sections were then labelled with multicolor fiducials (Tetraspecs) to facilitate high-precision correlation ^28,38^. Sections were imaged by 4-color widefield imaging and transmission electron microscopy or tomography.

To establish the specificity of the click-chemistry-derived fluorescent signal, we compared signal localization between samples of fixed cells grown on glass bottom 96-well plates with sectioned resin-embedded samples. In both cases, U2OS cells were loaded with a PC derivative bearing a palmitate at the *sn1* and the bifunctional fatty acid (Y) at the *sn2* position (PC(16:0|Y)) (Figure 1 A). After a 4 min lipid loading pulse, the probe was primarily localized at the plasma membrane (PM) and in endosomes for both sample types, confirming the specificity of the on-section click labelling (Figure 1 C, D). We validated the endosomal localization by co-staining with fluorescently labelled transferrin and LDL. The use of transferrin and LDL as suitable endosomal markers was demonstrated by co-staining with Rab5 (early endosomes), Rab7 (late endosomes), and Rab11 (recycling endosomes) (Extended Data 6). The lipid localization was further validated by assessing organelle ultrastructure using CLEM. The plasma membrane and endosomes were identified based on the fluorescent signal and imaged at high magnification by transmission electron microscopy (Figure 1 D). As expected, the plasma membrane and endosomal membranes exhibited high lipid fluorescent signals. Taken together, we conclude that the Lipid-CLEM workflow allows to assess the localization of near-native lipid probes within organelle membranes at the ultrastructural level.

### Polar resins are required for homogeneous fluorescence labeling

Measuring lipid densities at membrane nano domains requires information on the domains surface area and the lipid amount in the membrane domain of interest. 3D surface areas are obtained from electron-tomography data and relative lipid amounts from fluorescent signal intensities. Thus, to acquire accurate densities, lipid probes need to be evenly stained throughout the sections by on-section click chemistry.

To determine the homogeneity of the fluorescent signal throughout the resin, we labelled whole HM20 resin blocks derived from U2OS cells loaded with PC(16:0|Y). Resin blocks were then sectioned vertically and the sections imaged by fluorescence microscopy (Figure 2 A). HM20 samples showed signal confined to the edge of sections with a full width at the half maximum of 0.45 ± 0.10 µm (Figure 2 B, C). The resolution limit of the imaging setup was measured to 0.35 ± 0.03 µm using 100 nm fluorescent beads. The signal width of HM20 exceeds the resolution limit of the imaging setup. However, the section edge has a width of unknown dimension, which must be taken into consideration. Therefore, we additionally sectioned fully stained HM20 blocks horizontally into ultra-thin sections of 70 nm (Extended Data 7). The signal of the first four sections was found to be mutually exclusive, suggesting that the click labeling is limited to the section surface in HM20 samples.

**Figure 2.**
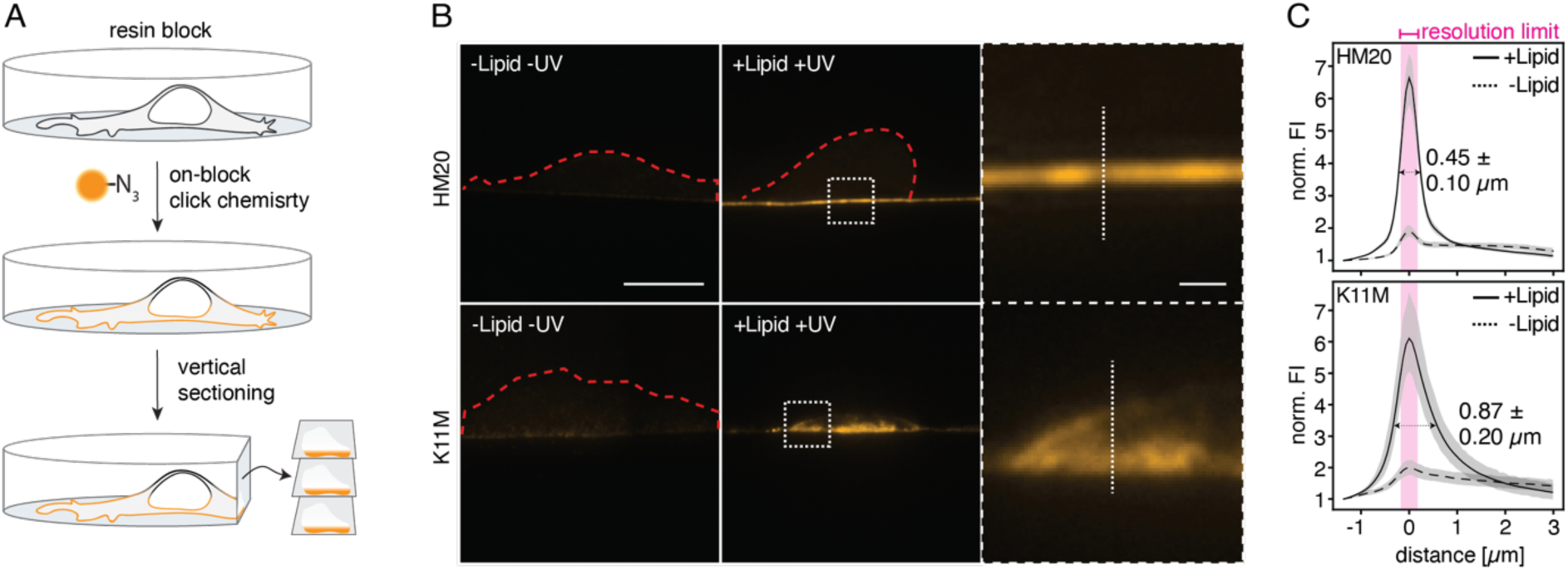
Comparison of dye diffusibility into HM20 and K11M resins. A: Workflow for testing the penetration depth of the click reaction mixture into resins. Whole resin blocks were stained and subsequently sectioned vertically to assess penetration depth. B: Representative images of stained and vertical sectioned blocks of HM20 (upper) and K11M (lower), of samples loaded with (+Lipid +UV) and without (-Lipid -UV) PC(16:0|Y). Scale bar of overview images: 10 µm; Scale bar of ROIs: 1 µm. Dotted red lines represent cell borders if not visible otherwise. C: The average line profile (indicated in B) of the fluorescence signal of vertical sections of HM20 and K11M with (black line) and without (dotted line) PC(16:0|Y). The error of the average line profile is shaded in grey and given as the confidence interval of 95%. The full width at half maximum (± standard deviation) for the lipid loaded samples is indicated. The purple bar indicates the resolution limit of the optical system as determined with 100 nm beads.

We hypothesized that the polar reactants of the click chemistry reaction mixture might not penetrate the hydrophobic HM20 matrix and therefore tested the polar K11M resin for fluorescent labeling. In K11M, the signal derived from vertically cut sections showed a significantly higher (students t-test, p-value: 1.82*10^-13^) average full width at half maximum of 0.87 ± 0.20 µm with visible internal membranes (Figure 2 B, C). The high penetration depth observed for K11M ensures the faithful assignment of lipid enrichment to membranes also in thicker sections used for tomography.

To ensure the specificity of the lipid signal in the K11M resin we next assessed the PC(16:0|Y) signal using CLEM. We observed the characteristic plasma membrane/ endosome localization of the lipid signal, indicating a specific staining (Figure 1 C, Figure 3 A, B). However, K11M samples exhibited some non-specific signal in nucleoli, a known click chemistry artifact ^39^, which has to be taken into account. One additional consideration to keep in mind is that membrane preservation in K11M is not as strong as in HM20 (Extended Data 8) ^40,41^.

**Figure 3.**
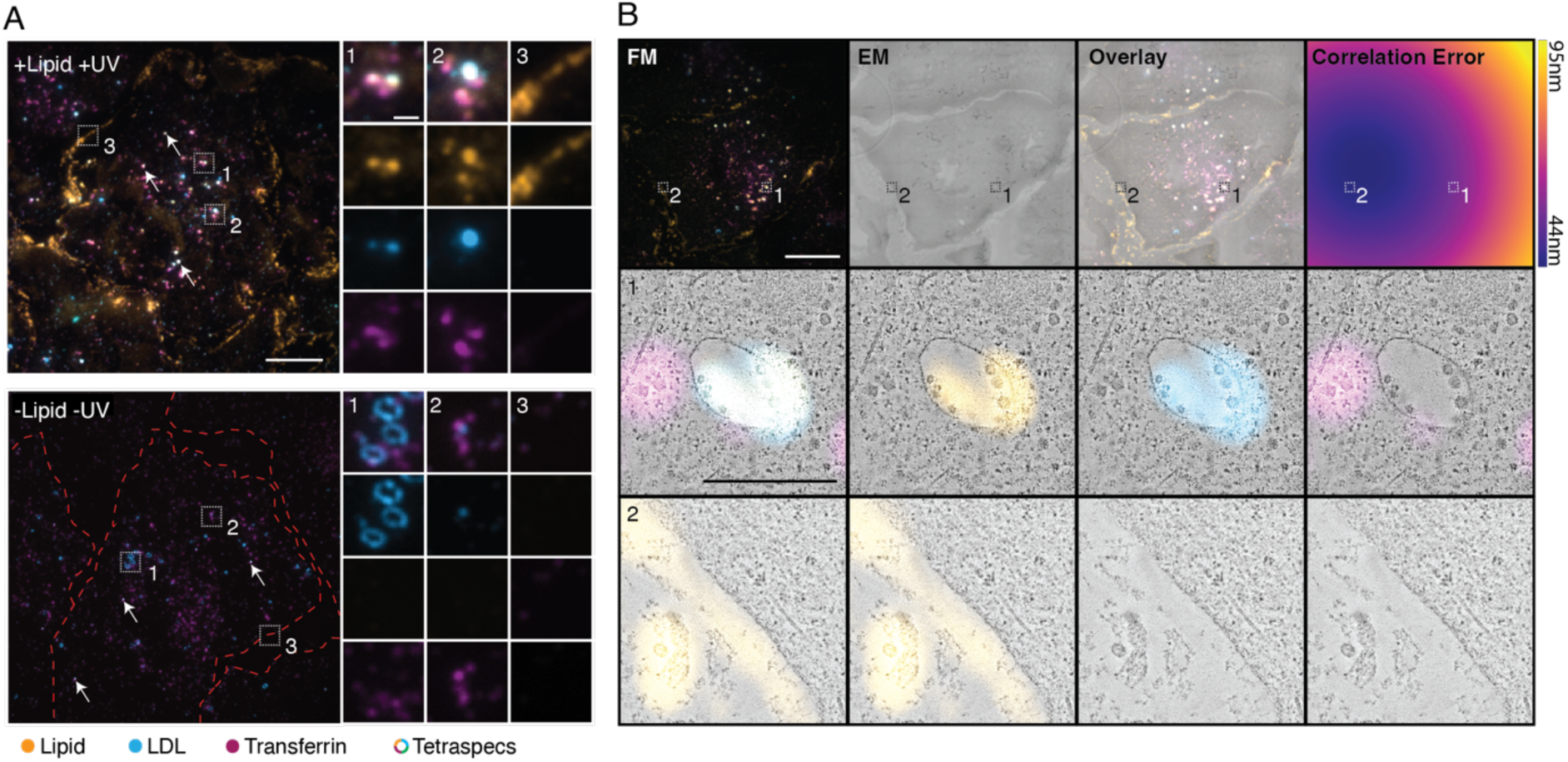
Lipid signal is well preserved and specific in K11M resin. A: U2OS wildtype cells were loaded for 4 min with PC(16:0|Y), UV crosslinked, high pressure frozen, embedded in K11M by automated freeze substitution and sectioned into 100 nm thick sections (+Lipid +UV). Negative controls were not UV activated and not loaded with PC(16:0|Y) (-Lipid -UV). All cells were co-labelled with fluorescent LDL (blue) and transferrin (magenta). Lipid signal is shown in orange. White dots correspond to multicolored Tetraspecs indicated by arrows. Dotted red lines represent cell borders if not visible otherwise. Scale bar of overview images: 10 µm; Scale bar of ROIs: 1 µm. B: The fluorescent overview image (FM), medium magnified TEM image (EM), overlay and correlation error map of 500 nm thick K11M sections are shown left to right in that order. A representative ROI for an endosome (1) and the plasma membrane (2) are shown, acquired at higher magnification by tomography. One image plane of the tomogram is shown. Scale bar of overview images: 10 µm; Scale bar of ROIs: 1 µm. Representative images of 3 biological replicates are shown.

We therefore propose two possible applications for on-section Lipid-CLEM, depending on the choice of resin used. Apolar resins such as HM20 are ideally suited to assign the localization of individual lipid species to well-preserved membrane ultrastructures at the surface of the resin. Polar resins such as K11M, on the other hand, are better suited for directly measuring lipid densities on 3D membrane structures.

### Sphingomyelin partitions into endosomal sub-compartments

We next analyzed lipid densities in the membrane compartments of the early endosome. Its complex membrane architecture features well-characterized protein domains, which are critical for protein sorting during membrane trafficking ^42,43^ and may play a role in lipid sorting as well ^34,35,44^. Partitioning of endosomal proteins into these long-lived compartments has been extensively studied and allows for straightforward classification. Here, we considered three endosomal compartments: the globular boundary membrane, intraluminal vesicles (ILVs) and recycling tubules. We chose to analyze the distribution of sphingomyelin in these endosomal compartments as an example of a typical plasma membrane and late secretory pathway lipid. To compare with bulk lipid density, we performed metabolic labelling with bifunctional palmitic acid, which is widely incorporated into the cellular lipidome.

Early endosomes were identified by the presence of both LDL and transferrin fluorescence signals, and high-resolution tomograms were acquired for the corresponding regions. All tomograms were reconstructed, and the set of endosomes which featured both intraluminal vesicles and recycling tubules, was used for further analysis. 3D membrane models were generated manually to determine the surface area of each compartment. The outlines of the 3D domains were z-projected into 2D images and blurred to generate a mask (Figure 4 A), to assign the fluorescent signals of sphingomyelin, LDL and transferrin to the compartments.

**Figure 4.**
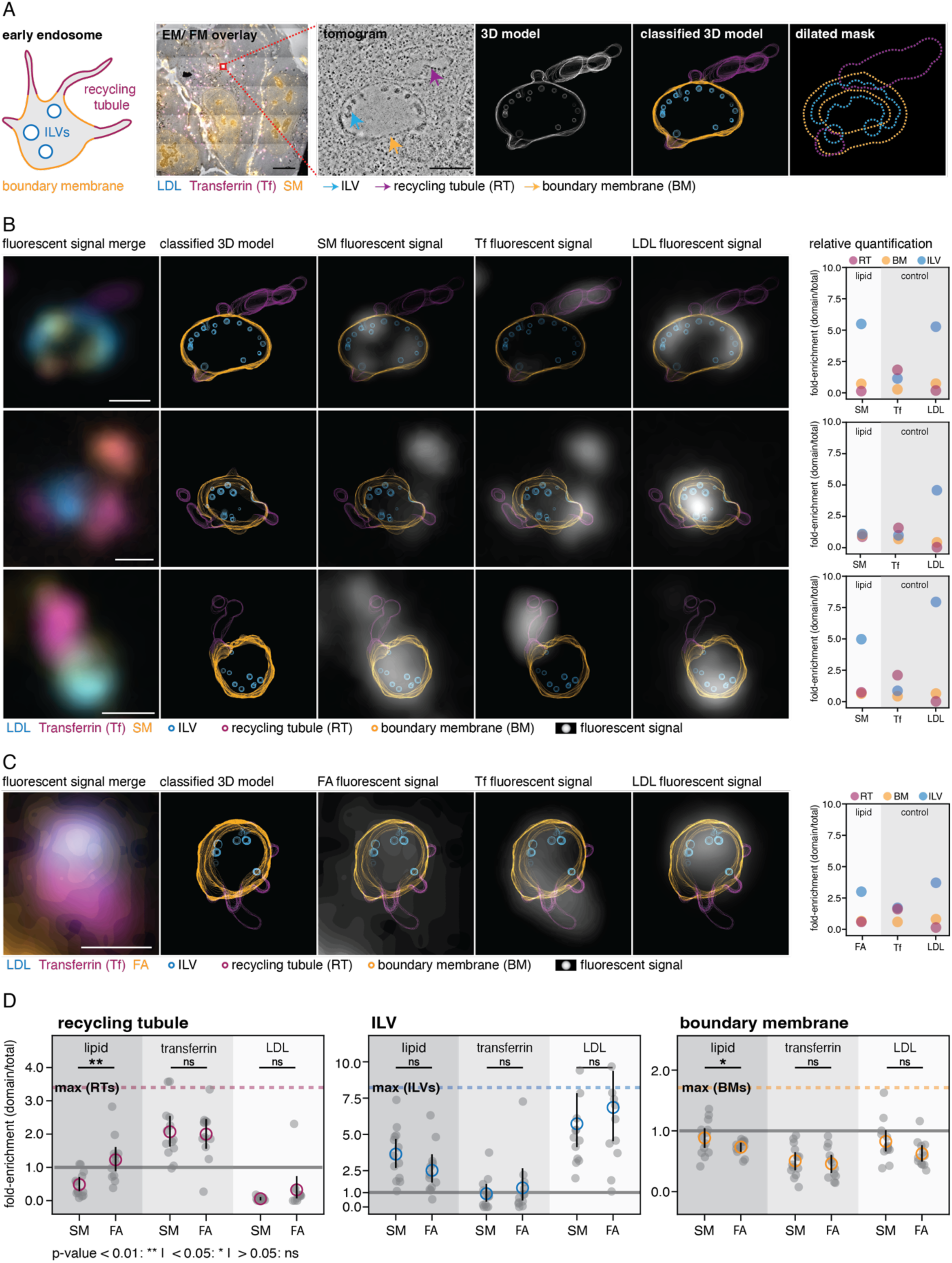
Sphingomyelin partitions differently into early endosome sub-compartments. A: CLEM data analysis: Images were acquired by fluorescent microscopy (FM) and electron microscopy (EM) at medium magnification. After correlation, the region of interest (ROI) was chosen based on the fluorescent signal of LDL (blue) and transferrin (magenta). Lipid signal is shown in orange. At the ROI a tomogram was acquired at high magnification. Tomograms were further analyzed if the endosomes contained boundary membrane (orange arrow), intraluminal vesicles (blue arrow), and a recycling tubule (magenta arrow). Based on the tomogram 3D models were derived. The z-projection of the 3D model outlines is shown. Membrane domains in the model are classified into boundary membrane (BM, orange), intraluminal vesicles (ILVs, blue), and recycling tubules (RTs, magenta) and the respective classes were blurred to create class-specific masks. Scale bar of overview images: 10 µm; Scale bar of ROIs: 500 nm. B: 3 representative endosomes are shown with the fluorescent overlay, classified 3D model, and model overlays with the fluorescent signal (grey) of sphingomyelin (SM), transferrin or LDL from left to right, respectively. Scale bars: 500 nm. Right panels: Relative densities of all fluorescent signals (SM, Tf and LDL) over all membrane classes: Recycling tubules (RT, magenta), intraluminal vesicles (ILV, blue) and the boundary membrane (BM, orange). C: Representative endosome of a cell treated with the bifunctional fatty acid and the corresponding quantification of the fluorescent signals. Scale bar: 500 nm. D: The relative fluorescent densities are plotted for all analyzed endosomes for SM and the metabolically labelled samples (FA). The mean densities are shown for the different domains: recycling tubule (magenta), boundary membrane (orange), and ILV (blue) and for the respective fluorescent signals of lipid, transferrin, and LDL. Maximum possible fold-enrichments are indicated by the dotted horizontal lines for the boundary membrane (orange), recycling tubule (magenta), and intraluminal vesicles (blue). No enrichment is indicated by the gray line at 1.0. Error bars are shown as the 95% confidence interval of the mean values. P-values are indicated and were determined using Student’s t-test. 2 biological replicates were performed both for SM and the fatty acid.

Fluorescent intensities were determined for each mask and normalized to the respective membrane domain surface area, resulting in fluorescence intensity densities. In regions where masks overlapped, partial pixel values were assigned to the respective masks based on probability density functions generated from the non-overlapping parts of the masks, when possible (see Materials and Methods for details). Finally, the fluorescent density of each membrane domain was divided by the fluorescent density of the whole endosome to determine relative enrichments (Figure 4 B, Extended Data 8). LDL and transferrin were used as internal standards for the analysis since they are enriched in intraluminal vesicles and recycling tubules, respectively ^31–33^. We observed the expected pattern of strong LDL enrichment in ILVs with a mean of 5.73 ± 0.99 relative density and strong transferrin enrichment in recycling tubules with a mean of 2.07 ± 0.23-fold density. These values can be considered as the practical upper boundaries of enrichment.

For sphingomyelin, we observed a slight depletion over the whole endosomal average in the boundary membrane with a mean relative density of 0.88 ± 0.08, enrichment in ILVs with a 3.64 ± 0.50 relative density and depletion in recycling tubules with a 0.49 ± 0.10 relative density. These results are indicative of differential localization and thus, sorting of sphingomyelin into different membrane domains in the early endosome (Figure 4 B, D; Extended Data 9). The respective values obtained for the metabolic labelling experiment with bifunctional palmitic acid were 0.73 ± 0.03 for the boundary membrane, 2.52 ± 0.48 for ILVs and 1.23 ± 0.19 for recycling tubules (Figure 4 C, D; Extended Data 10). Notably, the palmitic acid signal in endosomes was generally lower compared to other organelles, such as mitochondria and the ER, suggesting preferential incorporation into other organelle membranes. Taken together, lipid partitioning in endosomes was significantly different between a lipid population derived from metabolic labeling in comparison with a single sphingomyelin species and thus indicative of lipid sorting. Lipid sorting directly in the early endosome is further supported by the finding that SM and transferrin were taken up together by endocytosis, but separated in the early endosomal compartments, with transferrin being enriched in recycling tubule and SM in ILVs.

## Discussion

Here, we report a workflow for correlative light and electron microscopy to image lipids in cells (Lipid-CLEM) based on minimally modified lipid probes and on-section click chemistry labelling. Two distinct variants were developed for fluorescence labelling of (i) section surfaces and (ii) entire section volumes. Surface labelling in an apolar resin (HM20) is optimized for the preservation of ultrastructure and to precisely assign fluorescent signals in the z dimension. Full-volume labelling in a polar resin (K11M) allows for determining the densities of individual lipid species in complex membrane architectures in three dimensions. Our Lipid-CLEM approach for imaging individual lipid species in cellular membrane nanodomains complements existing methods for studying lipids at high resolution: Halogenated lipids have been visualized *in vitro* by single particle electron microscopy ^45^. Immunogold-labeling on sections of cells ^46,47^ allows for visualization of lipid class distributions by EM if suitable antibodies are available. Previous CLEM approaches based on minimally modified lipid probes have localized lipids or whole lipid classes qualitatively, however were typically not ideal for the preservation of membrane ultrastructure, or are limited to studying the outer leaflet of the plasma membrane ^12,26,27^.

We used Lipid-CLEM to address the question of lipid sorting in the early endosome and found that sphingomyelin is enriched in intraluminal vesicles and significantly depleted in recycling tubules. The observed depletion in recycling tubules cannot be explained by partitioning modes based on lipid asymmetry or sorting by curvature due to lipid shape ^48,49^, as these mechanisms can only account for much more moderate changes in partitioning. Since recycling tubules and the endosomal boundary membrane form one continuous membrane structure, the significant lower levels of sphingomyelin in one of the compartments (recycling tubules) suggests the existence of a diffusion barrier. The approximately 3-fold enrichment of sphingomyelin in intraluminal vesicles is most plausibly explained by either non-vesicular lipid transport in the lumen of the endosome or segregated sites of coordinated vesicle fusion and fission, which directly channel incoming lipid material to intraluminal vesicles. As a similar effect is observed during broad lipidome labelling, the latter hypothesis appears more likely. These findings allow us to draw the following conclusion: protein and lipid transport routes diverge at the early endosome as sphingomyelin and transferrin arrive simultaneously in clathrin-coated vesicles ^50,51^, but partition differentially into endosomal compartments.

Taken together, our methodology allows us to measure lipid densities in nanoscale compartments of biological membranes. By directly observing lipid partitioning into sub-compartments of the early endosome, we identify a checkpoint at which protein and lipid transport diverge during retrograde membrane trafficking. More generally, our approach will have major benefits for studying membrane nanoscale structures of complex architecture where a prominent role of lipids has long been hypothesized but was difficult to substantiate due to lacking methodology. Such domains include, for instance, signaling clusters at the plasma membrane, ER exit sites, nuclear pore complexes and cristae junctions in mitochondria. We anticipate that our Lipid-CLEM approach represents a major step forward towards defining membrane nanoscale architecture for both lipids and proteins. Data of this type are required for developing a broadly applicable consensus model of biological membranes.

## Acknowledgments

A.N. gratefully acknowledges financial support by the European Research Council (ERC) under the European Union’s Horizon 2020 research and innovation program (grant agreements no GA 758334 ASYMMEM and AURORA). AN acknowledges financial support by the Deutsche Forschungsgemeinschaft (DFG) via the TRR83 consortium. This research was supported by an Allen Distinguished Investigator Award, a Paul G. Allen Frontiers Group advised grant of the Paul G. Allen Family Foundation to AN. O.A. gratefully acknowledges financial support by the Israel Science Foundation (grant no. 3729/20), the European Research Council (ERC) under the European Union’s Horizon 2020 research and innovation program (grant agreement no 851080), the Henry Chanoch Krenter Institute for Biomedical Imaging and Genomics and support given by the Heineman Foundation through Minerva. O.A. is the incumbent of the Miriam Berman presidential development chair. We thank the following services and facilities at MPI-CBG Dresden for their support: the Electron Microscopy Facility, the Light Microscopy Facility, the Genome Engineering Facility, and the Scientific Computing Facility. We thank Jan Peychl, Britta Schroth-Diez, and Tobias Fürstenhaupt for their outstanding support and expert advice. We thank Paolo Ronchi and Alexander von Appen for expert advice during the development of the CLEM workflow.

## Author contributions

KB synthesized lipid probes. HML prepared samples and acquired the datasets. HML and WL sectioned blocks. HML and SK analyzed imaging data. FE constructed the LED photoreactors. HML, SK, NS, and MWB developed the CLEM lipid imaging workflow. HML, OA, and AN designed the project. OA and AN supervised research. HML and AN wrote the manuscript. All authors read and commented on the manuscript.

## Conflict of Interest Statement

The authors declare no conflict of interest.

## Data Availability

The imaging data can be accessed at https://doi.org/10.17617/3.LZPQ3K. The corresponding codes to analyze all data can be found at https://doi.org/10.5281/zenodo.14723745.

## Extended Data

**Extended Data 1:**
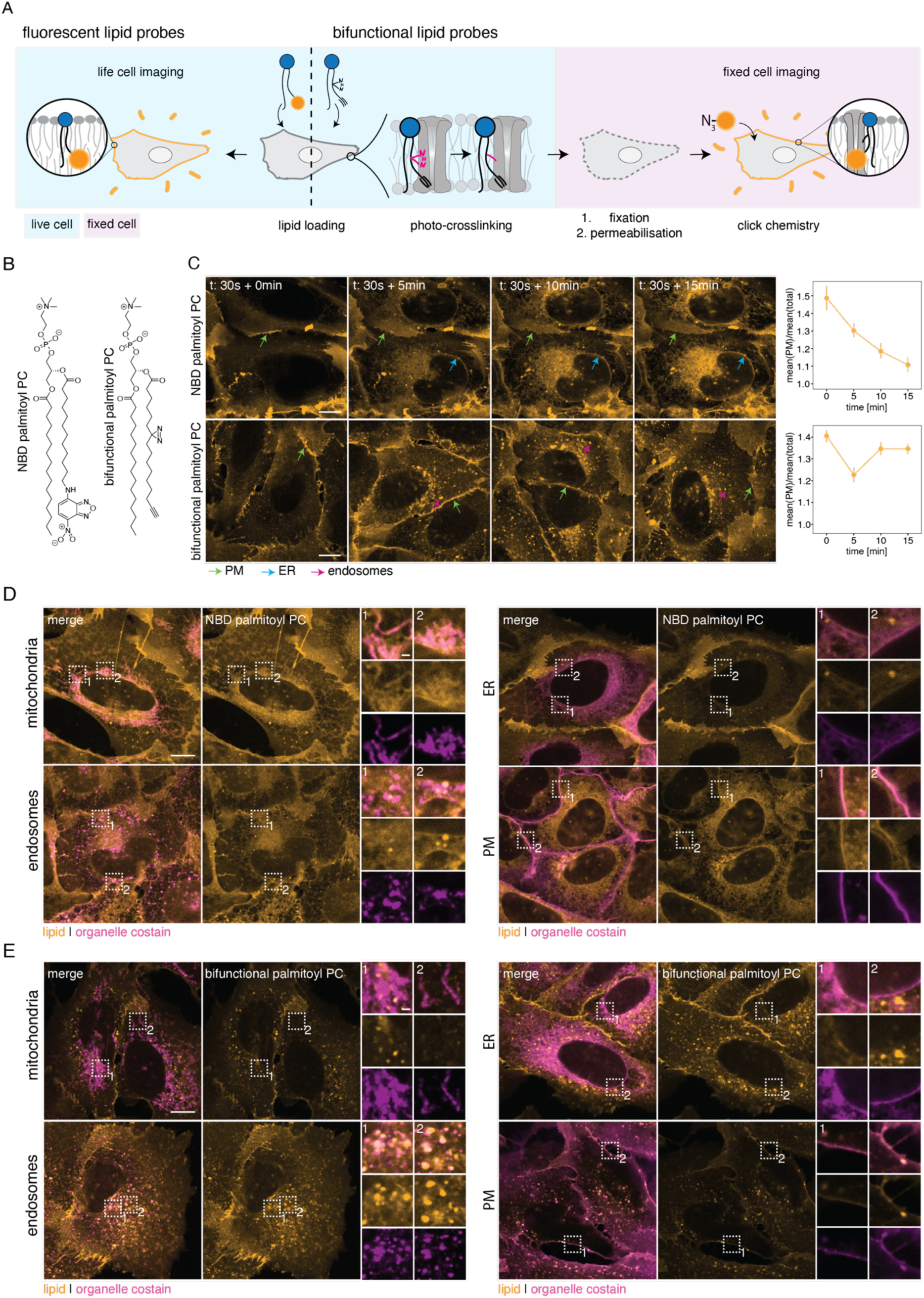
NBD and diazirine-alkyne labeled lipid probes show strong differences in organellar localization. A: Fluorescent lipid probes can be used for live cell imaging, while bifunctional lipid probes are suitable for fixed cell imaging. Bifunctional lipid probes require loading into live cells, photo-crosslinking, fixation, and labeling of the fixed samples by click chemistry. B: The chemical structures of the NBD PC(16:0 | 06:0-NBD) and bifunctional diazirine-alkyne PC(16:0|Y). C: U2OS wildtype cells were loaded with an NBD PC(16:0 | 06:0-NBD) for 30 s or a bifunctional diazirine-alkyne palmitoyl PC for 30 s and chased for 0, 5, 10 and 15 min each. Colored arrows indicate lipid localizations to different membrane-bound organelles. Lipid uptake was analyzed by assessing lipid amount in the plasma membrane over time relative to total lipid content. D, E: Lipid localization to specific organelles was confirmed with organelle-specific labels against mitochondria, endosomes, ER, and the plasma membrane for the NBD lipid and the bifunctional diazirine-alkyne lipid. Scale bar: 10 µm. Representative images of 3 biological replicates are shown.

**Extended Data 2:**
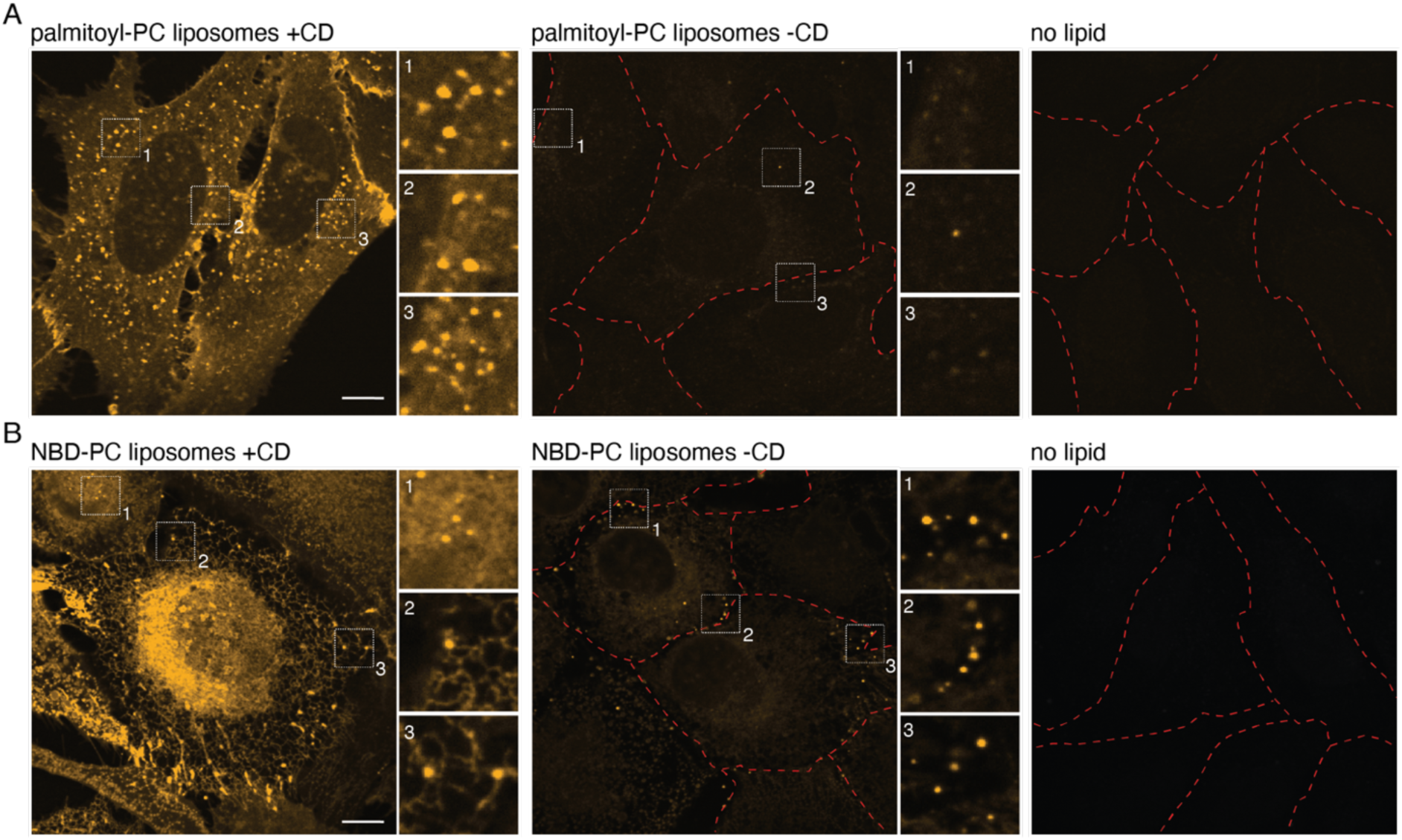
Bifunctional lipid probe-loaded liposomes are not taken up during lipid feeding. A: U2OS wildtype cells were treated with bifunctional lipid containing liposomes preincubated with alpha methyl cyclodextrin (+CD) or not (-CD). As a control samples without lipid treatment were also stained (no lipid). The samples without CD showed a low background, suggesting no uptake of whole liposomes by endocytosis. B: To ensure that the triton wash did not falsify this result, U2OS wildtype cells were also incubated with a mix of NBD-PC containing liposomes with and without CD during live cell imaging. Similarly low background was detected for loading conditions without CD. Scale bar: 10 µm.

**Extended Data 3:**
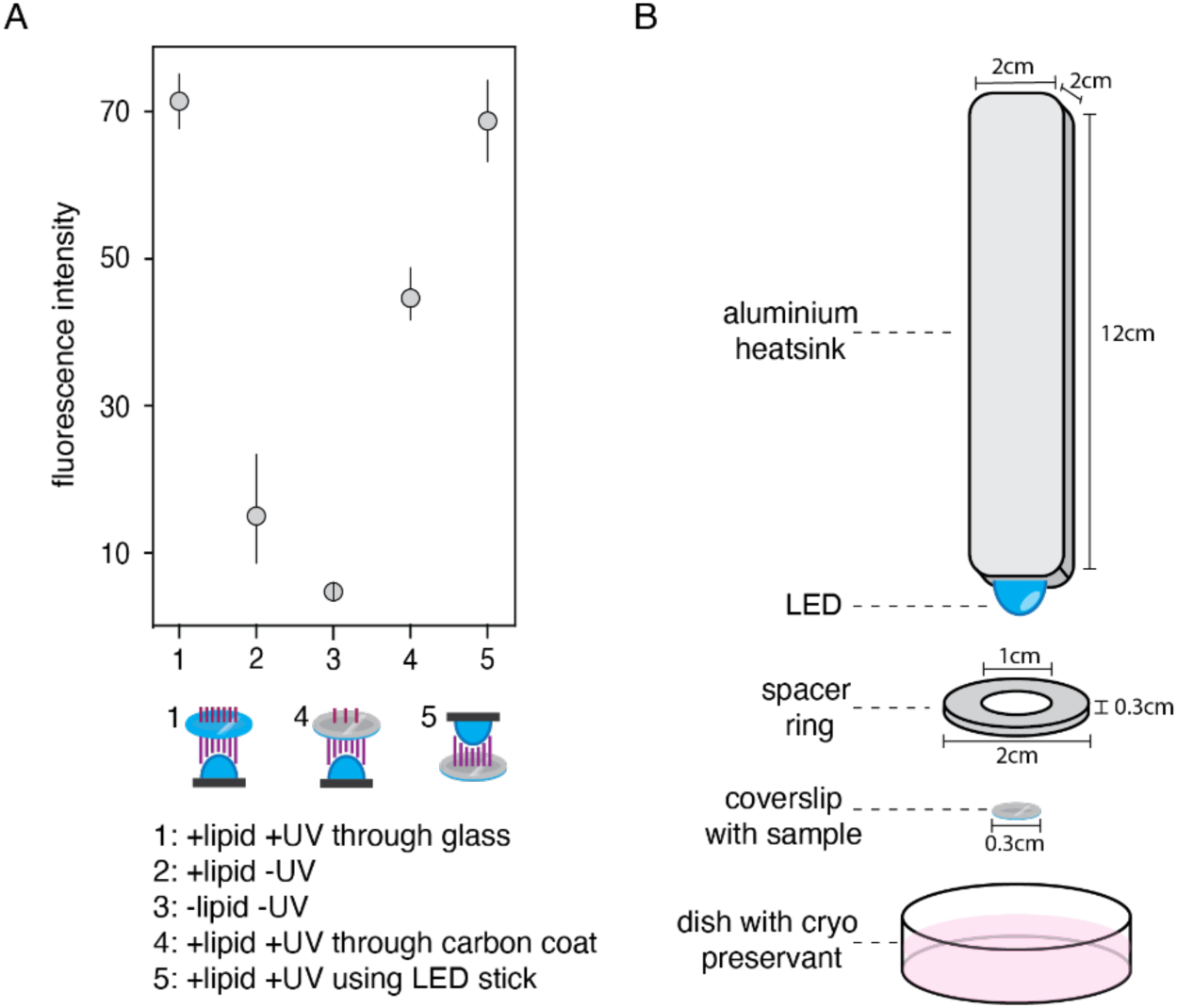
A water-resistant UV-LED setup allows for effective photo-crosslinking of carbon-coated samples. A: UV crosslinking efficiencies were measured in U2OS cells loaded with PC(16:0|Y). UV crosslinking was performed through glass (1), carbon coated glass (4) or from above (5). Negative controls were not treated with UV light (2) or PC(16:0|Y) (3). Mean values of fluorescent images are shown, error bars represent the standard deviation. The experiment was performed for 3 technical replicates. B: Schematic representation of the LED setup for illuminating samples in medium. Samples on coverslips are placed in a dish, a spacer ring is placed between the sample and the LED stick.

**Extended Data 4:**
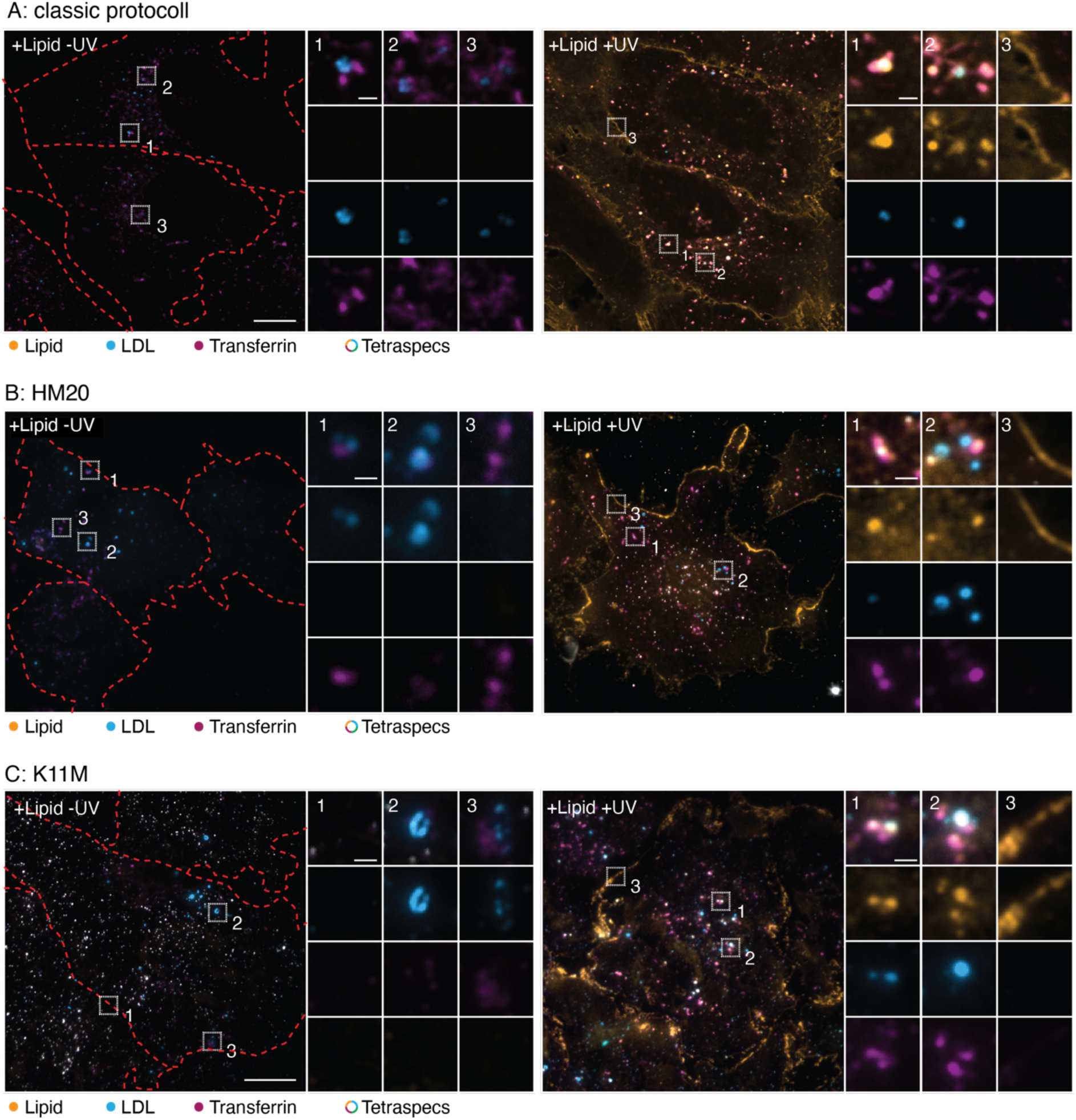
Non-photo-crosslinked bifunctional lipid probes are removed during automatic freeze substitution. A: U2OS cells were loaded for 4min with PC(16:0|Y), not photoactivated (+Lipid -UV) or photoactivated (+Lipid +UV), and prepared according to the “classic” lipid imaging protocol by chemical fixation, permeabilization, stain and imaging in PBS, or B: by embedding in HM20 and C: K11M according to the here proposed Lipid-CLEM workflow. The HM20 and K11M sections are 100 nm thin. Red-dotted lines show the cell outlines. Scale bars: 10 µm. The images of the right column are shown in Figures 1C and 2A.

**Extended Data 5:**
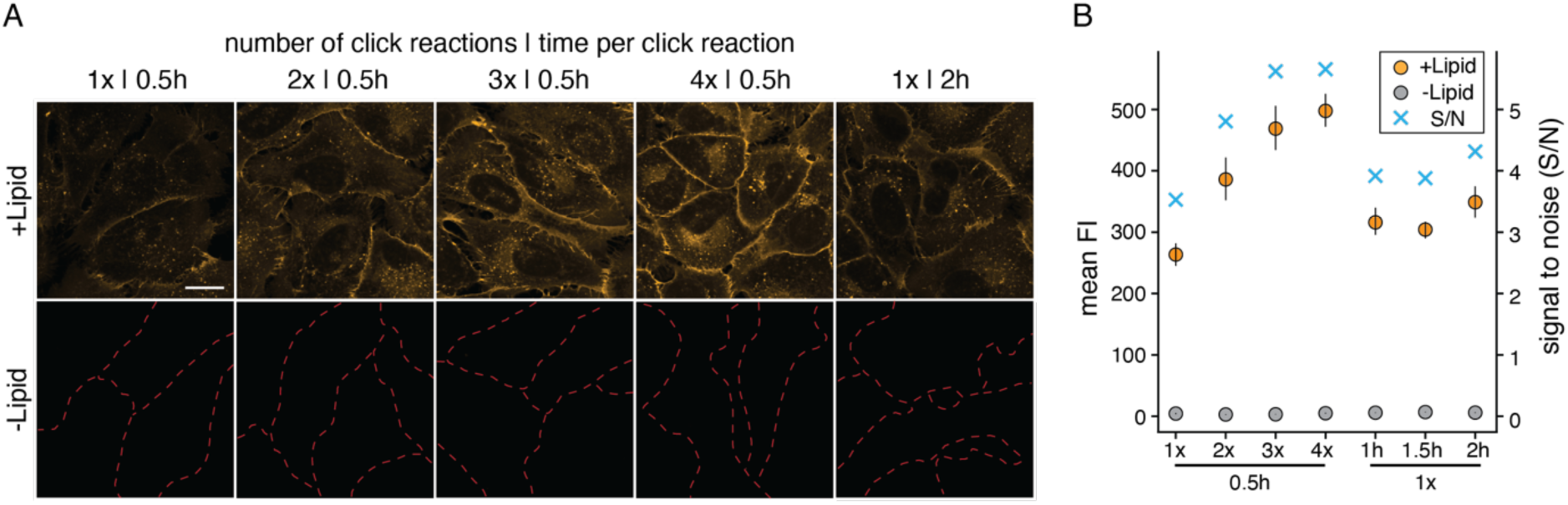
Repeated click labelling increases fluorescence signal. A: Representative fluorescence images of the optimization of the copper catalyzed click reaction are shown. U2OS wildtype cells were loaded for 4 min with PC(16:0|Y). Samples were stained 1 – 4 times for 30 min each, or stained once for the corresponding prolonged time. Negative controls (-Lipid) are shown in the bottom panels. Dotted red lines show the cellular outlines if not visible otherwise. Scale bar: 20 µm. B: Quantification of the experiment. Background corrected mean values of 3 technical replicates are shown. Error bars are shown as standard deviation. Lipid loaded samples are indicated in orange, negative controls in grey and the signal to noise ratio (S/N) as blue crosses.

**Extended Data 6:**
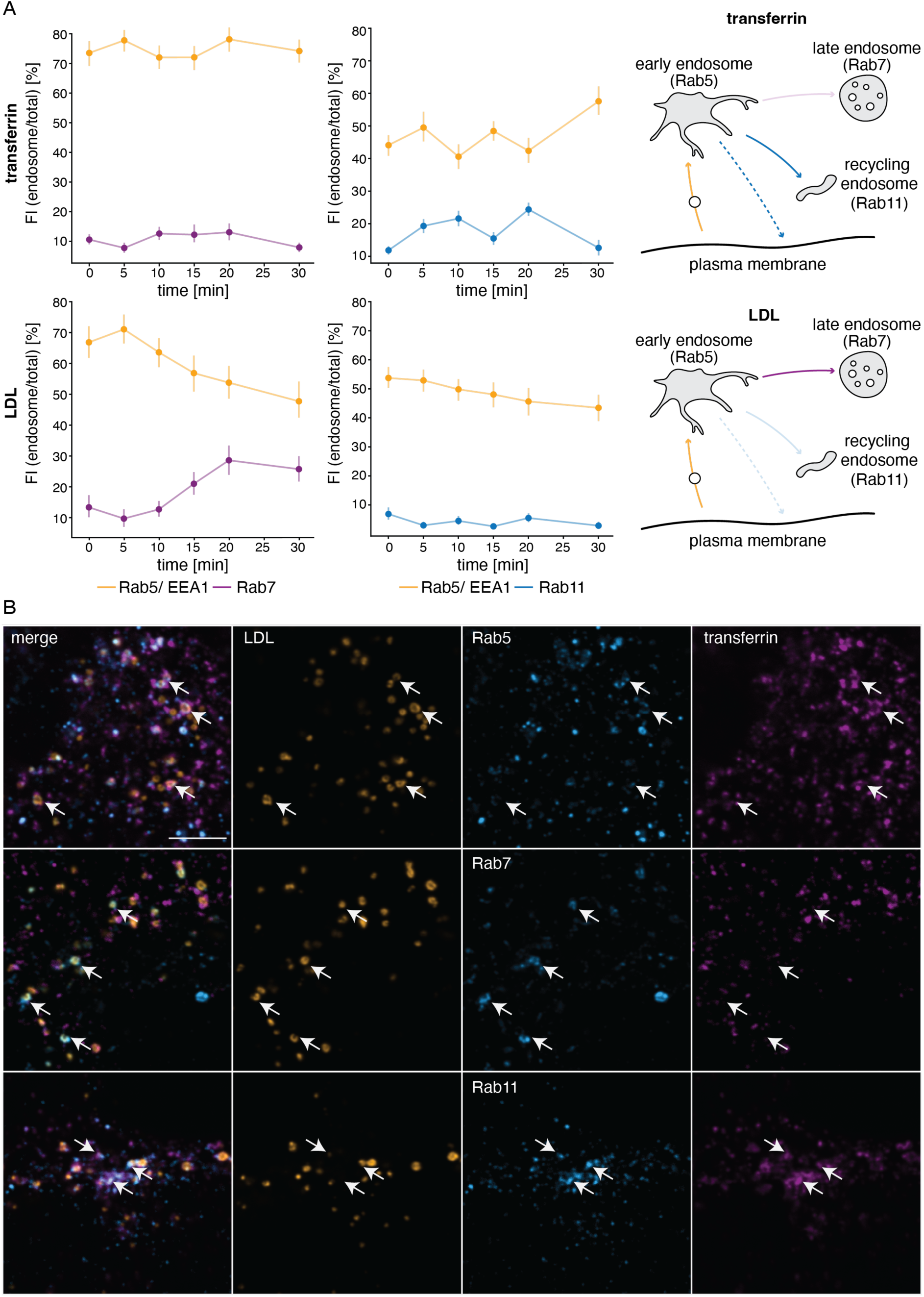
LDL and Transferrin specifically label endosomal compartments. A: Transferrin and LDL were first assessed separately for their localization to distinct endosomal compartments. Transferrin was loaded for 4 min, and chased for 0, 5, 10, 15, 20 and 30 min. LDL was loaded for 15 min and chased for 0, 5, 10, 15, 20 and 30 min. A mix of Rab5 and EEA1 was used to mark early endosomes, Rab7 was used for late endosomes and Rab11 was used for recycling endosomes. The fluorescent values of either transferrin or LDL were read out over the segmentation markers and normalized to total transferrin or LDL fluorescence signal. The analysis excludes compartments that are positive for both early and late endosomal markers or markers of early and recycling endosomes. Mean values are shown, error bars represent the standard deviation. Numbers above each mean value represent the counts of compartments in 1000s (k). The experiment was done as a single biological replicate. B: Transferrin and LDL were loaded together to assess their combined localization to distinct endosomal compartments. Representative images of U2OS cells labelled for 15 min with LDL (orange), chased for 15 min and labeled for 4 min with transferrin (magenta). Endosomal compartments were either immunolabelled with Rab5, Rab7 or Rab11 (blue). Scale bar is 5 µm.

**Extended Data 7:**
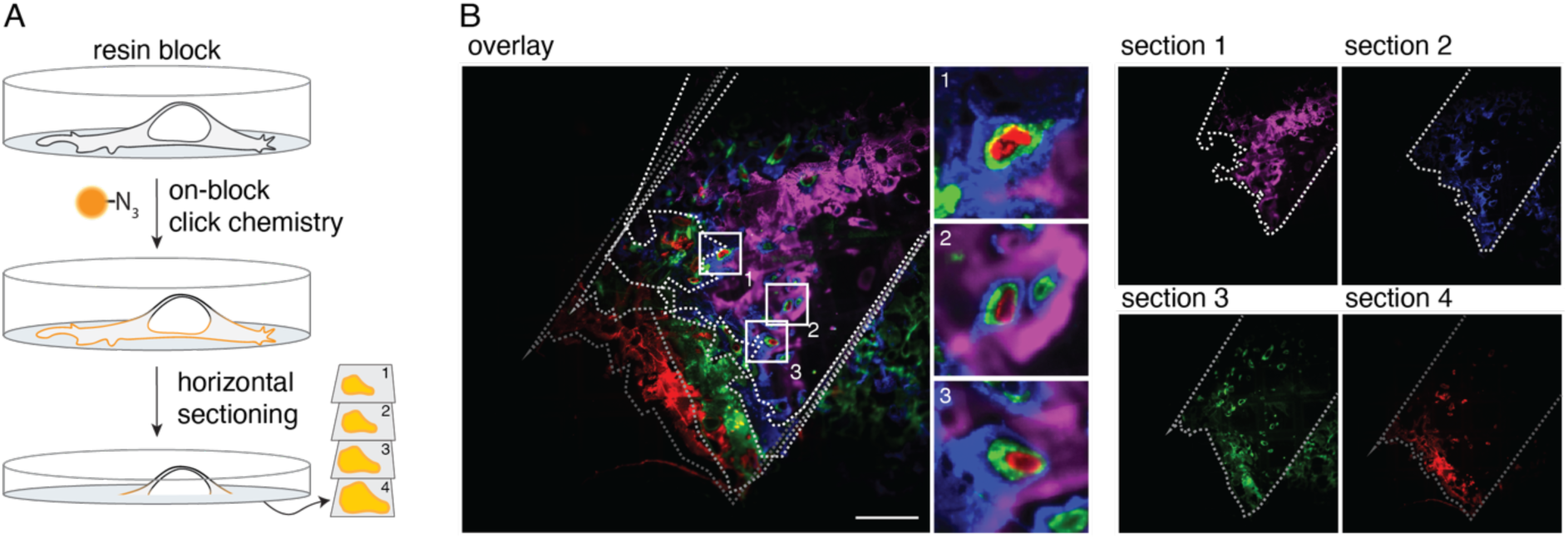
Penetration depth of the on-section click chemistry in HM20. A: A whole HM20 block was stained by click chemistry. The stained block was sectioned horizontally into 70 nm ultrathin sections. B: The fluorescence images of subsequent sections are shown. Different colors mark the section numbers. Magnified areas highlight the exclusivity of the stains between sections. White dotted lines outline the section borders. Scale bar: 100 µm. The experiment was performed as a single replicate.

**Extended Data 8:**
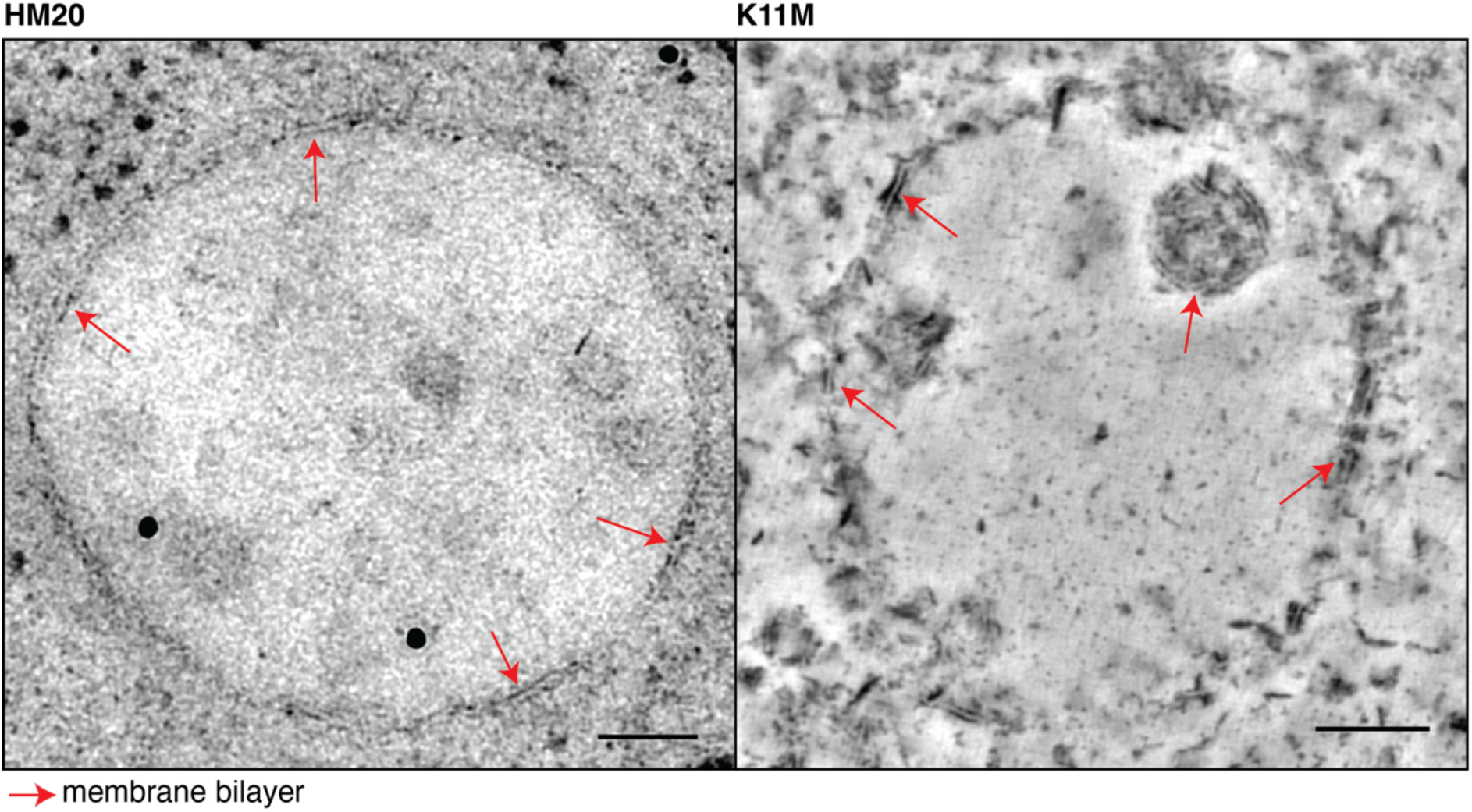
Preservation of membrane ultrastructure in different resins. On the left, an early endosome is shown embedded in HM20 (100 nm section). On the right, an early endosome is embedded in K11M (tomogram image from a 500 nm section). Red arrows point to regions where the lipid bilayer is visible. Scale bars are 100 nm.

**Extended Data 9:**
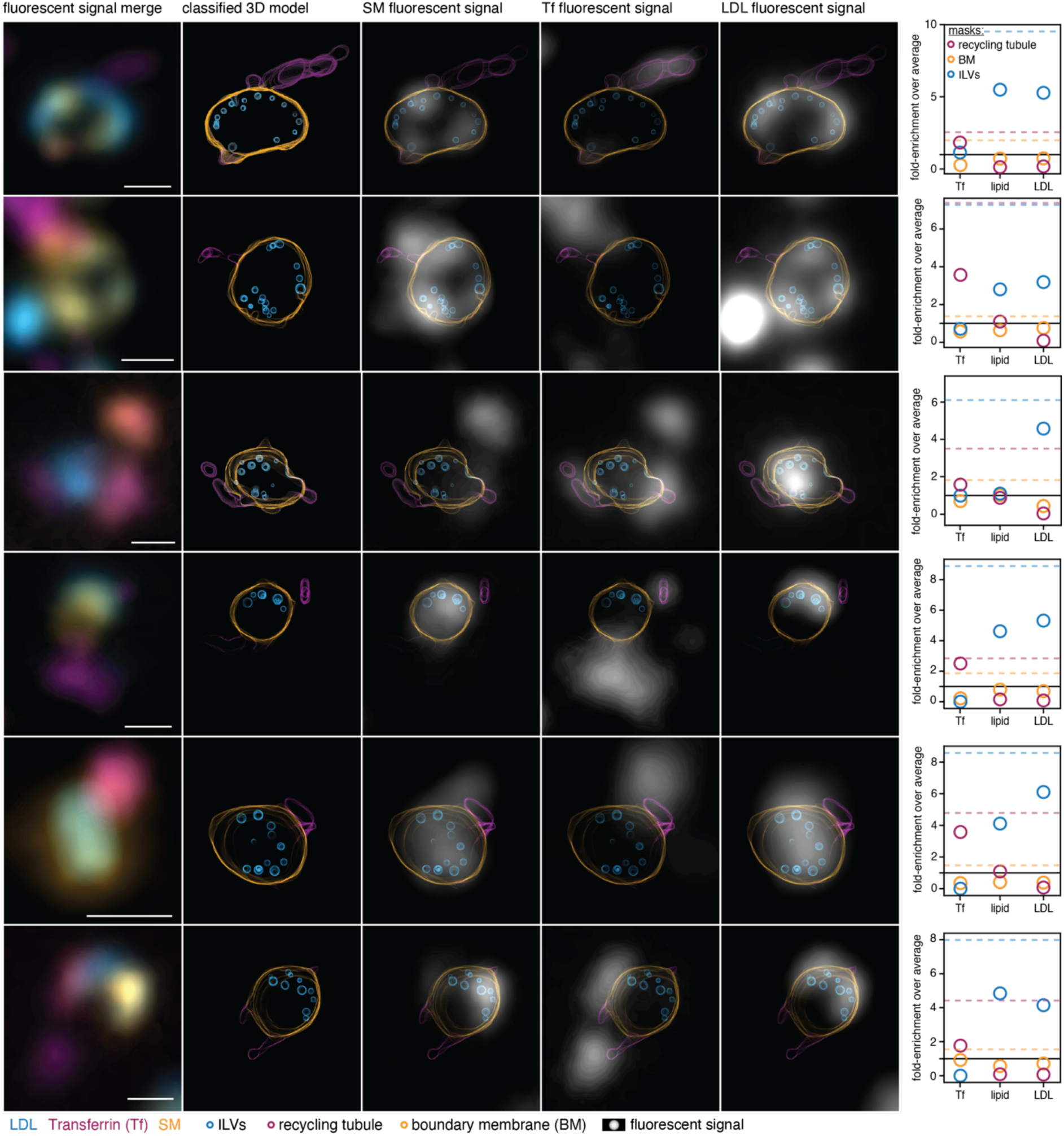
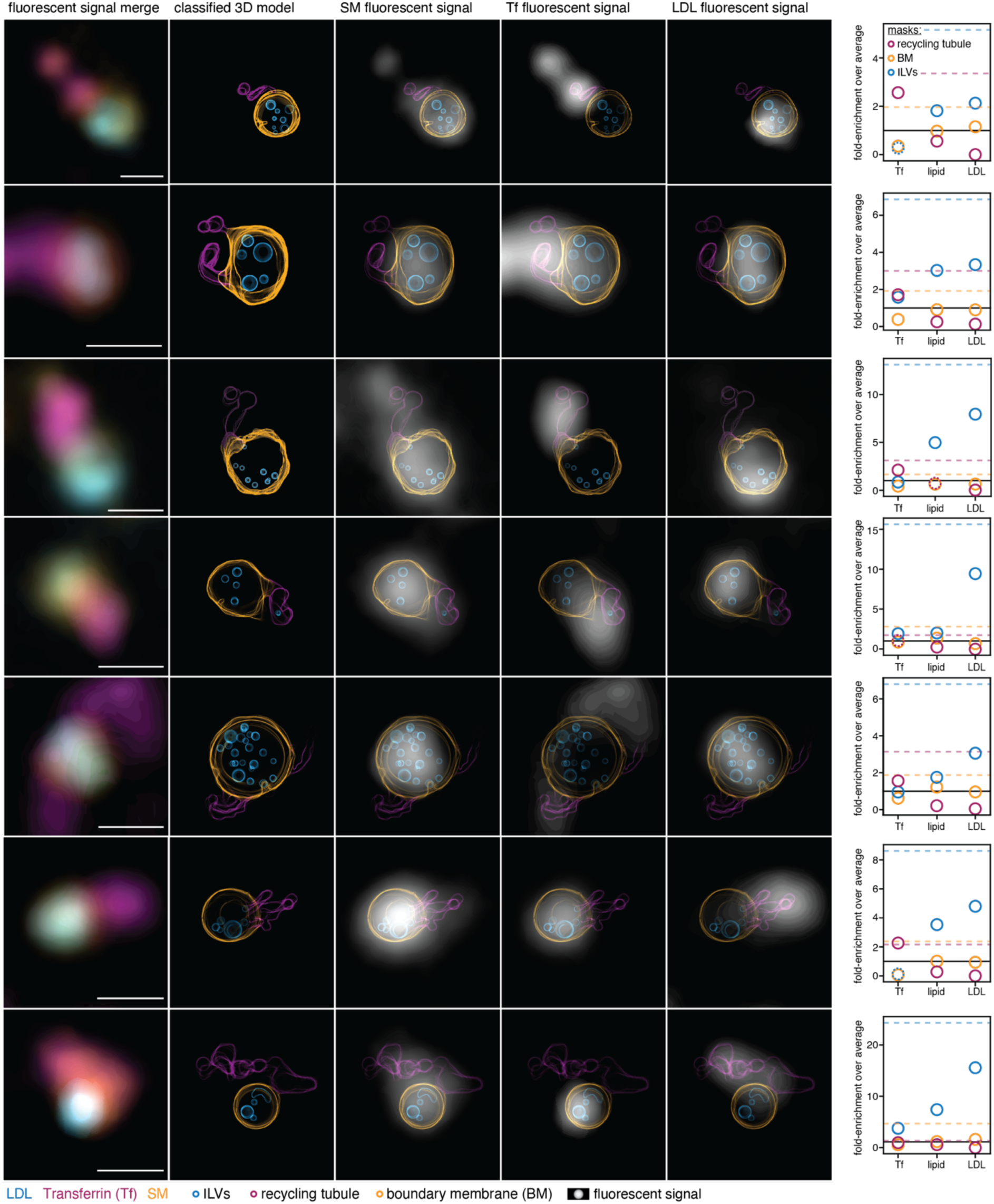
Analysis of early endosomes loaded with bifunctional sphingomyelin. All endosomes analyzed for the first biological replicate loaded with sphingomyelin (SM) are shown. The endosomes of row 1 and row 3 are shown also in the main text Figure 4. Column 1 shows the overlay of LDL (blue), transferrin (magenta) and SM (orange) fluorescence signals. Column 2 shows the max projected line profile of the classified 3D model. Column 3-5 show the model overlays with the fluorescent signal of SM, transferrin or LDL from left to right, respectively. On the right most side the enrichments of signal densities are depicted for all fluorescent signals over all membrane classes. Horizontal lines mark the maximum fluorescent density for recycling tubules (magenta dotted), intraluminal vesicles (blue dotted), the boundary membrane (orange dotted) as well as no enrichment (black). Scale bars: 500 nm. All endosomes analyzed for the second biological replicate loaded with sphingomyelin (SM) are shown. The endosomes of row 3 are shown also in the main text Figure 4. Column 1 shows the fluorescent overlay of LDL (blue), transferrin (magenta) and SM (orange). Column 2 shows the max projected line profile of the classified 3D model. Column 3-5 show the model overlays with the fluorescent signal of SM, transferrin or LDL from left to right, respectively. On the right most side the fold enrichments of signal densities are depicted for all fluorescent signals over all membrane classes. Horizontal lines mark the maximum fluorescent density for recycling tubules (magenta dotted), intraluminal vesicles (blue dotted), the boundary membrane (orange dotted) as well as no enrichment (black). Scale bars: 500 nm.

**Extended Data 10:**
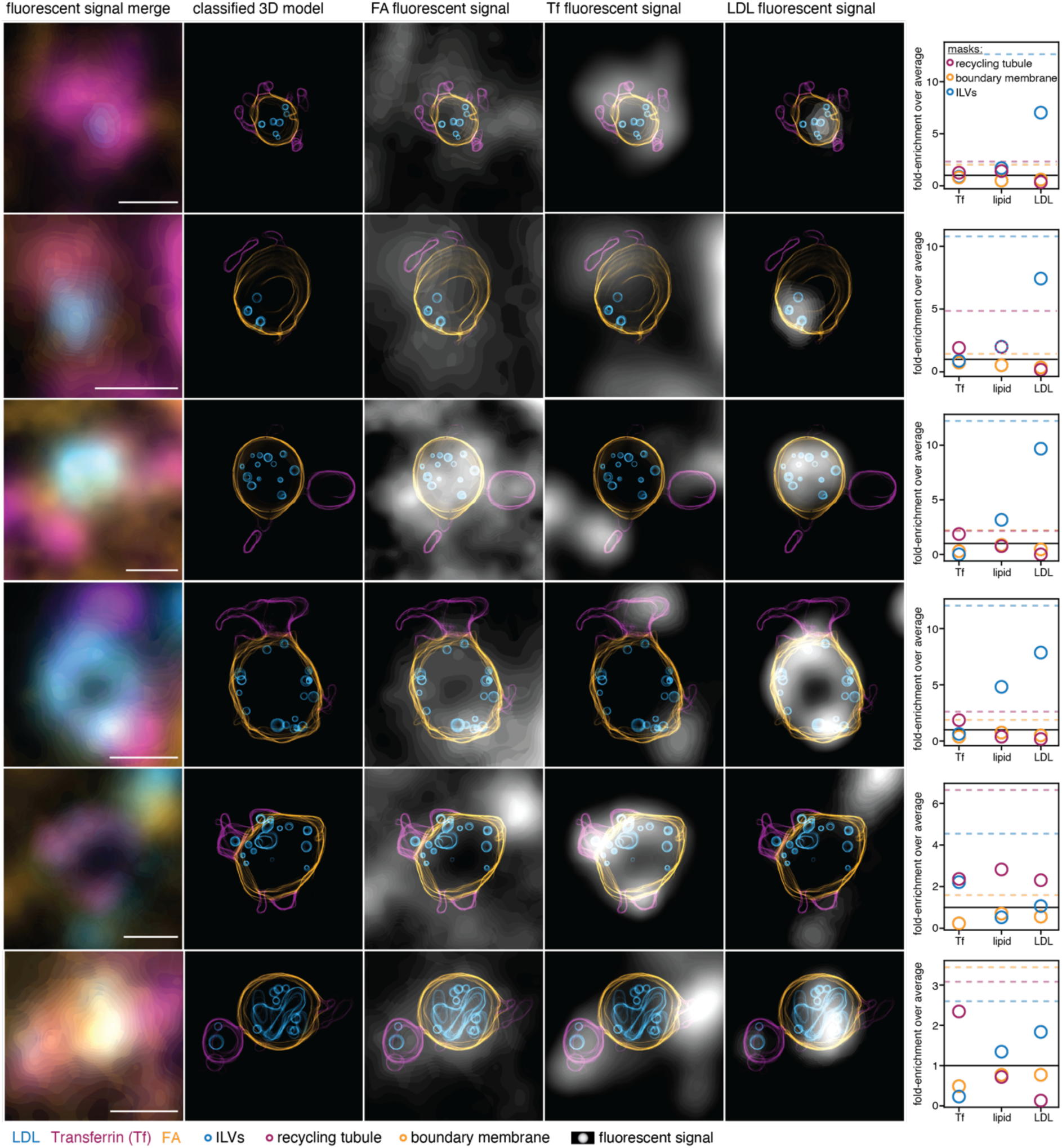
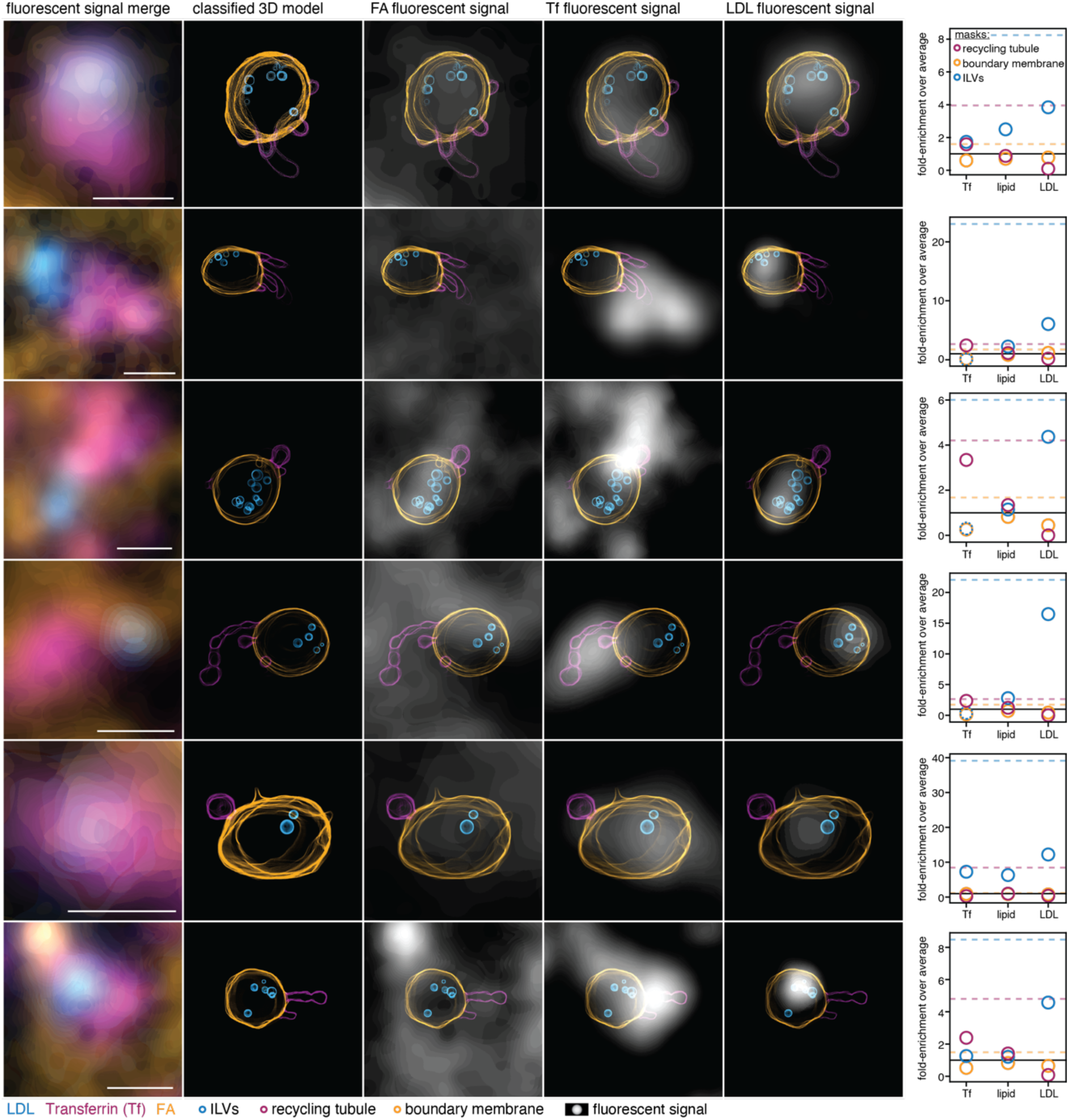
Analysis of early endosomes loaded with bifunctional fatty acid. All endosomes analyzed for the first biological replicate metabolically labelled with bifunctional palmitic acid (Y). Column 1 shows the fluorescent overlay of LDL (blue), transferrin (magenta) and SM (orange). Column 2 shows the max projected line profile of the classified 3D model. Column 3-5 show the model overlays with the fluorescent signal of SM, transferrin or LDL from left to right, respectively. On the right most side the fold enrichments of signal densities are depicted for all fluorescent signals over all membrane classes. Horizontal lines mark the maximum fluorescent density for recycling tubules (magenta dotted), intraluminal vesicles (blue dotted), the boundary membrane (orange dotted) as well as no enrichment (black). Scale bars: 500 nm. All endosomes analyzed for the second biological replicate metabolically labelled with bifunctional palmitic acid (Y). The endosome of row 1 is shown also in the main text Figure 4. Column 1 shows the fluorescent overlay of LDL (blue), transferrin (magenta) and SM (orange). Column 2 shows the max projected line profile of the classified 3D model. Column 3-5 show the model overlays with the fluorescent signal of SM, transferrin or LDL from left to right, respectively. On the right most side the fold enrichments of signal densities are depicted for all fluorescent signals over all membrane classes. Horizontal lines mark the maximum fluorescent density for recycling tubules (magenta dotted), intraluminal vesicles (blue dotted), the boundary membrane (orange dotted) as well as no enrichment (black). Scale bars: 500 nm.

## Notes

https://doi.org/10.17617/3.LZPQ3K

https://doi.org/10.5281/zenodo.14723745

### Competing Interest Statement

The authors have declared no competing interest.

