## Supplementary Information for "Visualizing sub-organellar lipid distribution using correlative light and electron microscopy"

### Materials and Methods

#### Sample preparation

##### Cell culture

U2OS wildtype cells (DSMZ, ACC785) were cultured in McCoy's 5A (1X) + GlutaMAX (gibco, #36600-021) medium supplemented with 10% fetal bovine serum (gibco, #10270-106) and 100 U/ml Penicillin-Steptomycin (gibco, #15140-122) at 37C with 5% CO2. Cells were passaged every 2-3 days using 0.05% Trypsin (gibco, #25300-054) up to a passage of 20. Cell seeding on carbon-coated disks is described below. Cells were seeded 24 h before starting the experiment into glass bottom 96-well plates (greiner, #655891) at a density of 9.000 cells per well.

##### U2OS Sec61 beta cell line

For live cell imaging with ER co-stain a Sec61 beta Halo U2OS cell line was made. The Sec61 beta Halo plasmid (#123285, addgene) was randomly integrated into the genome of U2OS wildtype cells (DSMZ, ACC785). 500 ng of the plasmid were transfected to 250.000 cells by neon electroporation at 1230 V, 10 ms for 4 pulses. Cells were expanded for 5 days, and stained using the Halo tag TMR (#G8251, Promega) according to the manufacture's recommendations. The cells were sorted using FACS. TMR-stained wildtype cells and non-stained transfected cells were used as negative controls.

##### Sapphire preparation and cell seeding

3 mm x 0.05 mm Sapphire disks (Wohlwend GmbH, #405 or Leica, #16702766 ) were coated with a total of approximately 15 nm carbon film using the safematic CCU-010 at 10 A current at $5.0^{-5}$ mbar pressure. Coated sapphires were baked overnight at 180 °C and used within one week. Immediately before cell seeding, sapphires were glow discharged at 15 mA negative, under 0.24 mbar for 90 s using a Pelco easiGlow (model 91000). Sapphires were placed in 35 mm glass bottom dishes (Cellvis, #D35-14-1.5-N) and pre-incubated with cell culture medium for 30 min at

37 °C. 240.000 U2OS cells per dish were seeded. Cells were grown for 12 – 18 h before high-pressure freezing.

#### Low-density lipoprotein labeling

Human low-density lipoprotein (LDL) (Invitrogen, #L3486) was fluorescently labeled with Alexa Fluor 488 carboxylic acid, succimidyl ester (Invitrogen, #A200000) in sodium bicarbonate buffer. Approximately 1 mg LDL was labeled covalently with 10 mg/ml (~200 nmol of reactive species) staining solution according to the manufacturer's recommendations for 1 h at room temperature. The freshly labeled mix was dialyzed against phosphate-buffered saline using a Slide-A-Lyzer Mini Dialysis (Thermo Scientific, # 88402) at 4 °C. The LDL solution in phosphate-buffered saline was supplemented with 10 % w/v sucrose and stored at -80 °C. Final LDL-AF488 concentrations were determined by a bicinchoninic acid assay.

#### LDL and transferrin loading

Prior to loading, cells were washed with serum-free medium. Labeled human LDL (see above) was dissolved in a serum-free medium at 5 µg/ml. Cells were incubated with freshly prepared LDL solution for 15 min and chased for another 15 min in serum-free medium without LDL at 37 °C. Labeled transferrin-AF647 (Invitrogen, #T23366) was dissolved in serum-free medium at 20 µg/ml, and cells were incubated with freshly prepared transferrin solution for 4 min at 37 °C.

#### Liposome preparation and lipid loading

The synthesis of all bifunctional lipids used in this study was previously reported [1]. A suspension of 1.5 mM bifunctional lipid, 0.75 mM 1-palmitoyl-2-oleolyl-glycerol-3-phosphocholine (Avanti, #26853-31-6), and 0.75 mM cholesterol (sigma, CAS: 57-88-5, #C8667-5G) in PBS was prepared. For NBD lipids (Avanti, #810131P, #810133P) a suspension of 1.5 mM NBD lipids, 0.75 mM 1-palmitoyl-2-oleolyl-glycerol-3-phosphocholine, and 0.75 mM cholesterol was prepared. Liposomes were made from this suspension using an Avanti Mini-Extruder with 0.1 µm polycarbonate membranes (Avanti, #610005-1Ea), by extruding for a minimum of 21 times. Liposomes and alpha-methyl-cyclodextrin (Bio-reagent, #CDexA-076/BR, CAS: 699020-02-5) were dissolved in serum-free medium to a final concentration of 0.5 mM and 4 mM respectively.

This mix was incubated for at least 30 min at 37 °C before loading on cells. Before lipid loading cells were gently washed 3 times with serum-free medium. Lipids were loaded for 4 min at 37 °C to cells unless stated otherwise. Cells were then washed with serum-free medium, followed immediately by UV activation. When loading bifunctional fatty acids for metabolic labeling the fatty acid was dissolved in 100 % ethanol and added directly in full medium to 5 µM for 17 h incubation duration.

#### UV irradiation

Samples were UV activated for 3 s using a custom built LED device (violumas, VC2X2C45L9-365) after lipid loading. The hand-held single LED was used for samples grown on carbon-coated disks and a custom multi-LED was used for samples in the 96-well plate format.

#### Chemical Fixation, Permeabilization and Stainings

Samples were fixed immediately after UV irradiation. For all control experiments unless stated otherwise cells were fixed using 4% PFA (CAS: 30525-89-4, TCI) in PBS (pH: 7.3) for 15 - 20 min at room temperature. For CLEM cells were fixed by high pressure freezing (see below). For conventional lipid imaging, fixed cells were washed 3 times with 100 mM glycine in PBS and permeabilized using 0.1 % Triton in PBS for 30 min at room temperature. Fixed and permeabilized samples were blocked at room temperature for 1 h in a blocking buffer consisting of PBS supplemented with 2% bovine serum albumin (BSA, Sigma, #A7030-10G). Primary Antibodies were incubated in blocking buffer for either 1 h at room temperature or overnight at 4 °C. Samples were washed 3 times for 15 min in blocking buffer at room temperature and incubated with the respective secondary antibody for 1 h at room temperature in blocking buffer. Samples were washed 3 times for 15 min in blocking buffer at room temperature and kept in PBS supplemented with 0.01 % sodium azide (Sigma Aldrich, #71290-100G). If samples were also treated with a copper-catalyzed click mixture as described below the antibody stain was performed first. Dyes for live cell imaging experiments were used according to the manufacturer. Table 1 shows all primary and secondary antibodies and dyes used with their respective dilutions.

Table 1| Dyes, primary and secondary antibodies

| Type | Target protein | Animal | Provider | Product umber | Dilution |
|---|---|---|---|---|---|
| prim | Rab5 | rabbit | Cell Signalling | 3547S | 1:500 |
| prim | Rab7 | rabbit | Cell Signalling | 9367S | 1:200 |
| prim | Calnexin 1 | mouse | abcam | ab112995 | 1:1000 |
| prim | Lamin A/C | mouse | Cell Signalling | 4777S | 1:1000 |
| prim | BAP31 | mouse | Enzo Life Sciences | ALX-804-601-C100 | 1:1000 |
| prim | Calreticulin | mouse | abcam | ab22683 | 1:1000 |
| prim | TOM20 | mouse | Santa Cruz | Sc-17764 | 1:1000 |
| prim | MAVS | mouse | Santa Cruz | Sc-166583 | 1:750 |
| dye | CellMask DeepRed | - | Invitrogen | C10046 | 1:3000 |
| dye | MemBrite Fix640/660 | - | Biotium | 30097-T | According to manufacturer |
| dye | MitoTracker Deep Red FM | - | Invitrogen | M22426 | 25 nM |
| dye | Janelia Fluor646 | - | Janelia | GA112A | 1:10000 |
| sec | AF647 anti rabbit | goat | Invitrogen | A21245 | 1:1000 |
| sec | AF488 anti mouse | goat | Invitrogen | A11001 | 1:1000 |

#### High-pressure freezing

High-pressure freezing (HPF) was performed using a Leica EM-ICE (Leica Microsystems). Before HPF, cells were treated as stated above with transferrin, LDL, and lipid. The sapphire discs are placed on the flat side of a 0.3 mm planchette (Wohlwend GmbH, #242) and covered using a 0.025/0.275 mm planchette (Wohlwend GmbH, #389) with the 0.025 mm side facing towards the cells. Planchettes were precoated with 1-Hexadecene (TCI, #H0610-100ML) and blotted before use. FluoroBrite DMEM (Thermofisher Scientific, #A1896701) supplemented with 20 % fetal bovine serum (gibco, #10270-106) was used as a cryo preservative.

#### Automatic freeze substitution

The protocol for the automatic freeze substitution (AFS) is depicted in Table 2. Frozen samples were transferred from liquid nitrogen to -90 °C cold acetone. Lowicryl HM20 (Embedding Kit 15924-1, Polysciences Inc.) and K11M (Embedding Kit 18163-1, Polysciences Inc.) were used according to manufacturer instructions. A Leica AFS2 machine and robot were used for embedding according to manufacturer instructions (Leica Microsystems). The samples were embedded in AFS sample wheels (Leica Microsystems, #16707154) and AFS sample ring holders (Leica Microsystems, #16707157). The freeze substitution medium contained 0.1% uranyl acetate (electron microscopy sciences, #22400) in acetone.

Table 2| Automatic Freeze Substitution Program.

| Step | $T_{start}$ (°C) | $T_{end}$(°C) | Slope | Time | Reagent | % | Transfer | Agitation | UV |
|---|---|---|---|---|---|---|---|---|---|
| 1 | -90 | -90 | 0 | 0:08 | FS medium | 100% | stay | on | |
| 2 | -90 | -90 | 0 | 2:00 | FS medium | 100% | stay | on | |
| 3 | -90 | -90 | 0 | 2:00 | FS medium | 100% | stay | on | |
| 4 | -90 | -90 | 0 | 2:00 | FS medium | 100% | stay | on | |
| 5 | -90 | -45 | 5 | 9:00 | FS medium | 100% | stay | off | |
| 6 | -45 | -45 | 0 | 7:00 | FS medium | 100% | stay | off | |
| 7 | -45 | -45 | 0 | 1:00 | Acetone | 100% | exch/fill | on | |
| 8 | -45 | -45 | 0 | 1:00 | Acetone | 100% | exch/fill | on | |
| 9 | -45 | -45 | 0 | 1:00 | Acetone | 100% | exch/fill | on | |
| 10 | -45 | -45 | 0 | 2:00 | HM20/ K11M | 10% | mix | on | |
| 11 | -45 | -45 | 0 | 2:00 | HM20/ K11M | 25% | mix | on | |
| 12 | -45 | -35 | 5 | 2:00 | HM20/ K11M | 50% | mix | on | |
| 13 | -35 | -25 | 2.5 | 4:00 | HM20/ K11M | 75% | mix | on | |
| 14 | -25 | -25 | 0 | 10:00 | HM20/ K11M | 100% | exch/fill | off | |
| 15 | -25 | -25 | 0 | 10:00 | HM20/ K11M | 100% | exch/fill | off | |
| 16 | -25 | -25 | 0 | 10:00 | HM20/ K11M | 100% | exch/fill | off | |
| 17 | -25 | -25 | 0 | 48:00 | HM20/ K11M | 100% | stay | off | X |
| 18 | -25 | 20 | 5 | 9:00 | HM20/ K11M | 100% | | off | X |
| 19 | 20 | 20 | 0 | 12:00 | HM20/ K11M | 100% | | off | X |

#### Sectioning

Fully cured resin blocks were sectioned using a Leica EM UC6 and a 35-degree diamond knife ultra (Diatome, #MT15630). Sections were cut to a thickness of 500 nm for tomography and 100 nm for standard TEM unless stated otherwise. HM20 was sectioned at a cutting speed of 0.8 mm/s and K11M at 1.4 mm/s. Sections were caught on copper 200-mesh copper grids with carbon support film (electron microscopy sciences, CF200-CU-50).

#### Click reactions

Chemically fixed and permeabilized samples in PBS and sections were stained using copper catalyzed click reaction by incubation at 37 °C with a click mix of 2 µM AF-594 picolyl-azide dye (jena bioscience, CLK-1296-1), 0.1 mM $CuSO_4$, 5 mM ascorbic acid and 0.5 mM THPTA (jena bioscience, CLK-1010-1G) in 100 mM HEPES at pH 7.3. Chemically fixed and permeabilized samples in PBS for spinning disc microscopy were stained once for 45 min. Sections were stained on grid mats (electron microscopy sciences, #71172) twice for 30 min at 37 °C and washed afterwards for 1 h at 37 °C in 100 mM HEPES and 10 times in deionized water by blotting at room temperature. All solutions were filtered using the Millex 0.22 µm filter (merck, #SLGP033RS).

##### Section post-processing

Sections were next stained before light microscopy on one side with multicolor TetraSpecs™ by incubation of grids for 5 min on a drop of freshly sonicated fiducials in PBS at 1:100 dilution for 100 nm sized TetraSpecs™ (ThermoFisher, # T7279) for K11M sections and 1: 25.000 dilution for custom ordered 50 nm TetraSpecs™ (ThermoFisher, # C47819) for HM20 sections. After imaging samples by light microscopy, grids were coated with freshly sonicated 15 nm gold beads on both sides at dilutions indicated by the supplier in PBS (CMC-Utrecht, #PAG 15NM/S). Following, K11M sections were stained with 1 % uranyl acetate solution (electron microscopy sciences, #22400) in deionized water for 7 min at room temperature, washed thoroughly in deionized water, and stained with a 0.04% (m/v) lead citrate (EMS, #17800) aqueous solution for 2 min at room temperature. Sections were thoroughly washed in deionized water and dried before imaging by electron microscopy.

#### Imaging

##### Spinning disk imaging

Images of chemically fixed and permeabilized samples in PBS were acquired on an Olympus IX83 microscope, equipped with a Yokogawa CSU-W1 SoRa unit, an ORCA-Fusion from Hamamatsu, and an ORCA-Flash 4.0 V3 digital CMOS camera using the FV10-ASW 1.7. software. A 100x immersion oil objective (Olympus UApoN OTIRF) at 65 nm pixel size was used for imaging. AlexaFluor488 (or NBD) was excited with a 488 nm laser and emitted light was collected between 500 and 550 nm. The AlexaFluor 594 azide was excited using a 561 nm laser and emitted light was collected between 580 and 654 nm. AlexaFluor 647 was excited with a 640 nm laser and emitted light was collected between 665 and 705 nm. Z-stacks were acquired with a 0.5 µm distance in z between frames. To ensure that all images were acquired at the same starting plane, the Olympus TruFocus Z-drift compensation system was used. All images for one dataset were acquired using identical settings.

#### Widefield imaging

CLEM sections were imaged by widefield imaging on an inverted stand Olympus IX83 by bypassing the disc. Whole grid overview images were taken with an Olympus U Plan SApo 20x (0.85 NA) Oil objective. Images were stitched using the Olympus cellSens 4.1 software. Image stacks of single grid squares were taken using an Olympus U-ApoN 100x (1.49 NA) Oil objective. Stack step size was chosen at 0.1 µm using a ZDC2 hardware autofocus and stage-top Z-piezo. The total stack size was 1 µm. All fluorophores were excited using a CoolLED pE-4000 at 100 % power for 500 ms exposure time. AlexaFluor(AF)405 was excited at 365 nm, using a U-FUNA (365/10 ET BP EX; DM 410; 440/40 ET BP EM) filter cube. AF488 was excited at 470 nm using a U-FBNA filter cube (Ex 482/25; DM 505; EM 5130/40). AF594 was excited at 550 nm, using a TxRed ET filter cube (560/40 ET BP EX; T585 LPXR DM; 630/75 ET BP EM). AF647 was excited at 635 nm using a Cy5 ET filter cube (620/60 ET BP EX; T660 LPXR DM; 700/75 ET BP EM). Detection was done with a Hamamatsu ORCA-Fusion BT Digital CMOS camera at a 23 MHz pixel clock.

#### Transmission electron microscopy imaging and Tilt-series acquisition

Transmission electron microscopy (TEM) on thin sections up to 100 nm in thickness was done on a Tecnai T12 (Thermo Fisher Scientific, Hillsboro, Oregon, USA) at 100 kV acceleration voltage with the standard single tilt sample holder. The images were taken on an F416 camera (Tietz Video and Image Processing Systems GmbH, Gilching, Germany) at 4096 x 4096 pixels. Tomography on thick sections from 500 nm was done on a Tecnai TF30 G2 FEG-TEM (Thermo Fisher Scientific) at 300 kV acceleration voltage with a model 2040 Dual-Axis Tomography Holder (Fischione Instruments, Export, PA, USA). The images were captured on a Gatan OneView camera (Gatan, Pleasanton, California, USA) at 4096 x 4096 pixels. Imaging on both microscopes was done with the program SerialEM [2]. Grids were placed into the sample holder with the sample facing down. Images at the T12 were acquired at magnifications of 2900 x (3.761 nm pixel size) for grid square overview images and 13000 x (0.8211 nm pixel size) for high magnification zoom-ins. Images at the TF30 were acquired at magnifications of 3100 x (3.897 nm pixel size) for the grid square overview and 12000 x (1.0308 nm pixel size) for the tomogram acquisition. Tilt series were acquired from two axes, from 0° to -60° and 0° to 60°.

#### Image processing, correlation, and tomogram reconstruction

Image and data post-processing was performed using python [3] , Ilastik [4], Prism, ICY/ ec-CLEM [5], etomo, imod/ 3dmod [6], and ImageJ/ Fiji [7,8]. The python version 3.8, ImageJ/ Fiji version 2.14.0/1.54f, Ilastik version 1.4.0-OSX, imod/3dmod version 4.11.25 and ICY (first version) with the ec-CLEM plugin were used.

##### Ilastik models

Ilastik was used for segmentation for the following experiments: Segmentation of the plasma membrane of the live cell (uptake traces of NBD lipid probes) samples and fixed samples (uptake traces of bifunctional lipid probes, uptake traces of transferrin and LDL). In all cases, we used the 2-stage Autocontext pixel classification workflow of Ilastik. The training was performed in 2D and using different frames of the Z-stack.

##### Correlation and tomogram reconstruction

Fluorescent image stacks were acquired by widefield fluorescence imaging. Images were first preprocessed. Out-of-focus images in each stack were removed. The image stacks were merged using ImageJ/ Fiji with the average intensity setting. The contrast and brightness were adjusted per each merged image. To account for shifts due to chromatic aberration ICY ecCLEM was used for shift correction by picking the center of single TetraSpecs. The lipid channels were kept as the original unchanged image. For correlation between the fluorescence image and the medium magnification TEM image, the ICY ecCLEM plugin was used. Based on the fluorescent signal different regions for tomography were chosen. Tomograms were reconstructed using etomo.

#### CLEM penetration depth

Line profiles were taken of in focus single plane images (line width 10) in the lipid channel using ImageJ/ Fiji, perpendicular to the section surface. Line profiles were taken only in areas with cells (determined with the respective signal from LDL and transferrin). Profiles were saved as .csv files. Per biological replicate (2 per condition) 15 line profiles were taken. To determine the resolution of the microscope setup 100 nm fiducials were analyzed similarly but using a line width of 1.

#### Analysis of fluorescence signal densities on membrane structures from CLEM data

Image and data post-processing was performed using python [3] (version 3.8). In the following, "channel" is defined as the fluorescence signal detected for transferrin, LDL, or lipid, respectively, and "mask" indicates the segmentation masks for the outer globular membrane of the endosome, intraluminal vesicles, and recycling tubules. 3D models of the membranes of endosomes were generated manually using imod/ 3dmod. The step size in z was adjusted to the reported thickness retrieved after the tomogram reconstruction of each tomogram. Membranes were classified into the main globular membrane (outer membrane), recycling tubule (membranes protruding outward from boundary membrane, meshed with cap), and intraluminal vesicles (vesicles enclosed by the outer membrane and/ or recycling tubule, meshed with cap). Surface areas were determined using *imodinfo*. The outlines of each object were generated using .wimp files of the separated objects and the python script *suman_wimp_lineprofiles.py*. Using the python script *ML_pdf_analysis.py* line profiles were blurred to generate masks. The width of the Gaussian blur (sigma: 30) was set to capture the majority of fluorescence based on the transferrin and LDL channel. Masks were split into regions with no overlap with the other masks, overlap with another mask, or overlap with all three masks. The background corrected fluorescence intensity of each channel (LDL, transferrin, lipid) was read out over each mask region without overlap. For the pixel values for non-overlapping masks, a probability density function was generated using a Gaussian kernel with a bandwidth of 1.0 using the *scikit-learn kernel density estimator*. Using the probability density function per mask and channel, partial pixel values were assigned in regions of overlapping masks at the determined probability. The fluorescent pixel values were recombined per mask from the non-overlapping region and regions with overlap. The total fluorescence intensity was determined per mask and channel and divided by the membrane surface area of the corresponding mask to obtain the fluorescence density of each mask and channel. The background corrected fluorescence per channel was also read out over the total mask (combining the 3 masks of the outer membrane, intraluminal vesicles, and recycling tubule) and normalized over the total membrane surface of the whole endosome. To determine fold-enrichments over the average per channel, the fluorescence density assigned to each mask was divided by the fluorescence density of the whole endosome. Maximum possible fold-enrichments were calculated by taking the ratio of the total membrane surface area over the membrane surface area of the mask.

#### Calculation of fluorescent density, relative density, and maximum possible densities

For every fluorescent marker, the fluorescence intensity was measured per mask and normalized to the respective membrane domain surface area, resulting in fluorescence intensities per membrane $nm^2$ (fluorescence densities):

$\text{fluorescence density(domain)} = \frac{fluorescence\ intensity\ (in\ mask\ of\ domain)}{surface\ area\ (domain)}$, with the domains of the outer membrane, intraluminal vesicles, or recycling tubules. To calculate the relative fluorescence density, the fluorescent densities of the membrane domains were each normalized over the mean fluorescent density of the full endosome:

$$\text{relative fluorescence density} = \text{fluorescence density (domain)}/\left[\frac{fluorescence\ intensity\ (total\ mask)}{surface\ area\ (total)}\right]$$

To determine the maximum possible enrichment in a domain, we assume that the total fluorescence intensity of a label is localized to one domain. Maximum fold enrichments are calculated as follows: $\max fold\ enrichment = \left[\frac{total\ fluorescence\ intensity}{surface\ area\ of\ domain}\right] / \left[\frac{total\ fluorescence\ intensity}{total\ surface\ area}\right]$, and thus: $\max fold\ enrichment = \left[\frac{total\ surface\ area}{surface\ area\ of\ domain}\right]$.
